# A high-resolution mRNA expression time course of embryonic development in zebrafish

**DOI:** 10.1101/107631

**Authors:** Richard J. White, John E. Collins, Ian M. Sealy, Neha Wali, Christopher M. Dooley, Zsofia Digby, Derek L. Stemple, Daniel N. Murphy, Thibaut Hourlier, Anja Füllgrabe, Matthew P. Davis, Anton J. Enright, Elisabeth M. Busch-Nentwich

## Abstract

We have produced an mRNA expression time course of zebrafish development across 18 time points from 1-cell to 5 days post-fertilisation sampling individual and pools of embryos. Using poly(A) pulldown stranded RNA-seq and a 3′ end transcript counting method we characterise the temporal expression profiles of 23,642 genes. We identify temporal and functional transcript co-variance that associates 5,024 unnamed genes with distinct developmental time points. Specifically, a class of over 100 previously uncharacterised zinc finger domain containing genes, located on the long arm of chromosome 4, is expressed in a sharp peak during zygotic genome activation. The data reveal complex and widespread differential use of exons and previously unidentified 3′ ends across development, new primary microRNA transcripts and temporal divergence of gene paralogues generated in the teleost genome duplication. To make this dataset a useful baseline reference, the data are accessible to browse and download at Expression Atlas and Ensembl.

## Introduction

Gene regulatory interactions are the fundamental basis of embryonic development and transcription is one of the major processes by which these interactions are mediated. A time-resolved comprehensive analysis of relative mRNA expression levels is an important step towards understanding the regulatory processes governing embryonic development. Here we present a systematic assessment of the temporal transcriptional events during this critical period in zebrafish (*Danio rerio*).

The zebrafish is a unique vertebrate model system as it possesses high morphological and genomic conservation with humans, but also experimental tractability of embryogenesis otherwise only found in invertebrate model organisms such as *Drosophila melanogaster* or *Caenorhabditis elegans*. Features such as very large numbers of offspring, *ex vivo* development and embryonic translucency have enabled comprehensive forward and reverse genetic screens (Amsterdam et al., 1999; Driever et al., 1996; Haffter et al., 1996; Kettleborough et al., 2013; Moens et al., 2008; Varshney et al., 2015) as well as high-throughput drug discovery approaches (Murphey et al., 2006; North et al., 2007; Peterson et al., 2000; Peterson et al., 2004; Stern et al., 2005). Together with a high quality genome (Howe et al., 2013b) only comparable in vertebrates to mouse and human, this has led to many important discoveries in areas such as zygotic genome activation (ZGA)(Lee et al., 2013), blood stem cell biology (Bertrand et al., 2010; Kissa and Herbomel, 2010), but also highly translational findings directly affecting human health (Li et al., 2015; Tobin et al., 2010).

The morphological processes underlying the transformation of a fertilised egg into a free swimming fish have been studied extensively owing to the ease with which embryogenesis can be observed and manipulated (Behrndt et al., 2012). This has identified many genes that drive crucial steps of the differentiation process, however the wealth of morphological phenotype data has not been matched with a systematic analysis of the corresponding molecular events. Baseline transcriptomic datasets in other species have greatly improved knowledge of relative levels of gene expression as well as alternative splicing events (Boeck et al., 2016; Brown et al., 2014; Gerstein et al., 2014; Graveley et al., 2011; Hashimshony et al., 2015; Klepikova et al., 2016; Owens et al., 2016; Tan et al., 2013).

Previous baseline transcriptome work in zebrafish has focused on either certain aspects of development such as the maternal-zygotic transition (Aanes et al., 2011; Harvey et al., 2013; Lee et al., 2013), the identification of specific transcript types (Pauli et al., 2012) or promoters (Gehrig et al., 2009; Nepal et al., 2013), on gene annotation improvement (Collins et al., 2012) or have a limited number of time points and replicates (Yang et al., 2013).

Using poly(A) pulldown RNA-seq and a 3′ end enrichment method called DeTCT (Collins et al., 2015) we have generated a comprehensive polyadenylated RNA expression profile of normal zebrafish development. We have taken advantage of the large numbers of offspring and have divided the 6 day period of embryonic development into 18 stages. For each stage we have processed up to 29 biological replicates consisting of single or pooled embryos.

We have placed particular emphasis on making the presented data accessible without the need for computational processing. The data are available to browse and download in a user-friendly interface at Expression Atlas and as selectable tracks in the Ensembl genome browser. Furthermore, incorporation into the next Ensembl gene set update will significantly improve zebrafish gene and transcript isoform annotation.

This developmental polyadenylated RNA baseline reference enhances our understanding of the gene regulatory events during vertebrate development and provides a time-resolved transcriptional basis for the comprehensive annotation of functional elements in the zebrafish genome using existing and future genome-wide datasets (Haberle et al., 2014; Kaaij et al., 2016; Lindeman et al., 2010; Potok et al., 2013; Vastenhouw et al., 2010).

## Results

### A high-resolution transcriptional profile of zebrafish development

A total of 18 developmental time points were sampled in this study (Figure 1A). Four time points are before or at the onset of zygotic transcription, four during gastrulation, three during somitogenesis, three during prim stages – 24, 30 and 36 hours post-fertilisation (hpf) – and every 24 h from 2 to 5 days post-fertilisation (dpf). This gives detailed coverage of all the important developmental processes taking place during this time period. At each time point, samples were collected for both RNA-seq and DeTCT (RNA-seq, pools of 12 embryos, 5 replicates; DeTCT, pools of 8 embryos, 11-12 replicates and individual embryos, 11-12 replicates). We obtained an average of 3.8 million reads per sample for the RNA-seq and 9 million reads per sample for the DeTCT data (Figure 1B). The number of genes detected rises steadily as development proceeds and a wider range of genes are expressed (Figure 1C, Supplemental Figure 1-1). With a cut-off of greater than or equal to 5 normalised counts it rises from a mean of 10,395 to 18,225 for the RNA-seq (6,259 to 15,215 in the DeTCT data).

For quality control purposes, counts from the RNA-seq data, normalised for library size, were first used to produce a sample correlation matrix (Figure 1D). This shows the excellent agreement among biological replicates and close relation of neighbouring stages in terms of gene expression profiles. To give a high level overview of the data, principal component analysis (PCA) was used to visualise patterns. These show close clustering of biological replicates and a smooth transition from one stage to another through developmental time. The first six principal components (PCs) explain just over 90% of the variance in the data (Figure 1E-G and Supplemental Figure 1-2); the first component accounts for 58.2% of the observed variation. Summaries of the expression patterns underlying the PCs are shown in Figure 1H. Broadly, the genes that contribute the most to the first PC are either low at early time points and increase towards the end of the time course or the inverse (high early and decreasing later). PC2 represents genes with a peak at gastrula and somite stages. Genes contributing to PC3 show two peaks of expression (onset of transcription and days 3-5). The other patterns are more complex (Supplemental Figure 1-2) although PC6 shows a clear spike of expression starting at the 1000-cell stage (3 hpf), which coincides with the start of zygotic transcription.

In order for this dataset to be as useful as possible to the scientific community, we have made it available as sequence with metadata through the European Read Archive and in multiple locations for viewing. Pre-computed count profiles (FPKM) are available to search/browse at Expression Atlas (http://www.ebi.ac.uk/gxa/experiments/E-ERAD-475)(Petryszak et al., 2016), allowing comparison of the expression profiles of multiple genes across the entire time course (Figure 2A). This is searchable by gene/allele name, Ensembl/ZFIN ID and GO terms. It is also possible to start with a single seed gene and have Expression Atlas add in similarly expressed genes (as assessed by k-means clustering) for comparison (https://www.ebi.ac.uk/gxa/FAQ.html#similarExpression). The data are also viewable in Ensembl as separate stage-specific selectable tracks along with one merged track (http://www.ensembl.org/Danio_rerio). The aligned reads can be displayed as either coverage graphs or read pairs. Reads that span introns can be viewed to investigate alternative splicing in a stage-specific manner (Figure 2B-C).

**Figure 1.**
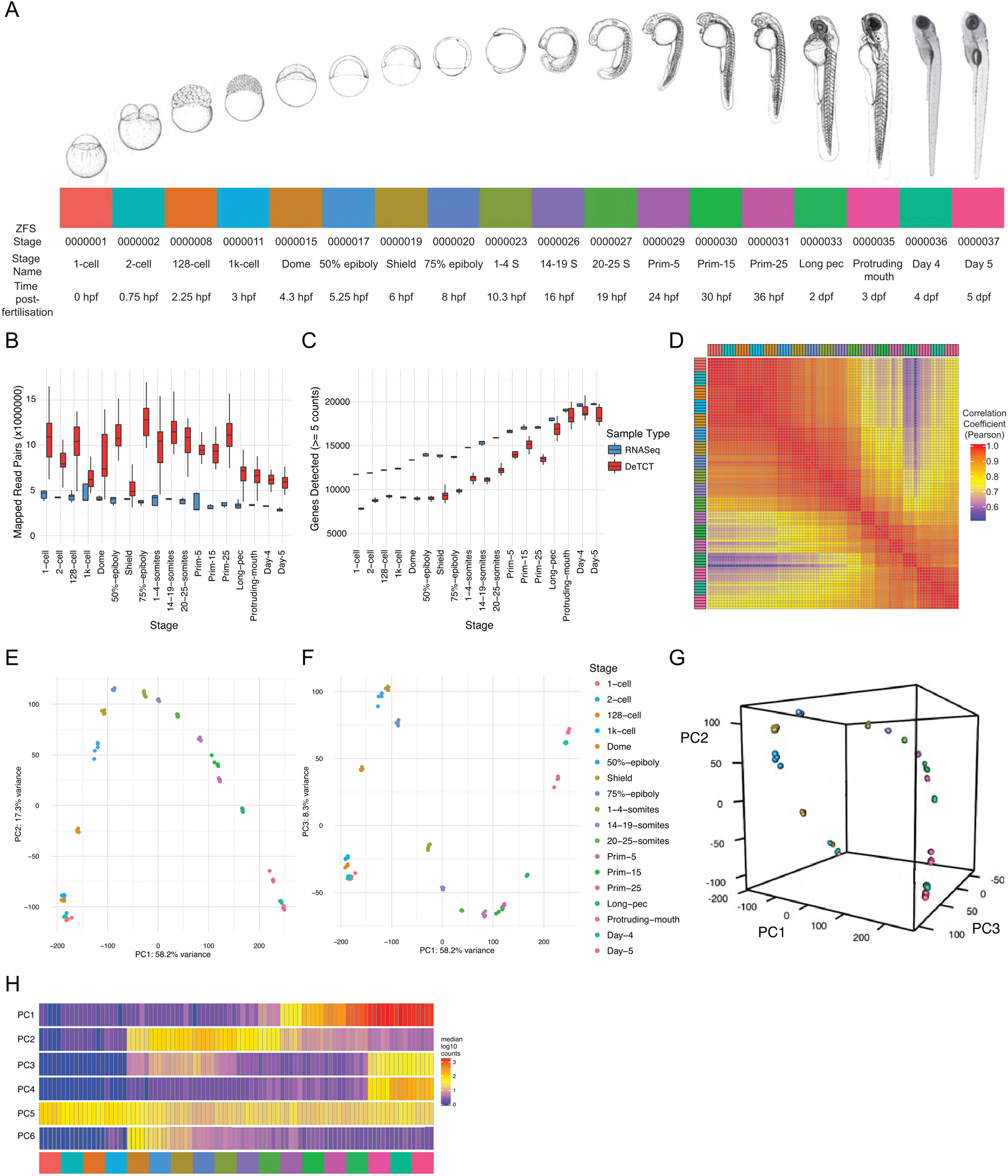
A transcriptional map of development. (A) Stages represented in this study. Images of embryos from 1-cell to protruding mouth are from (Kimmel et al., 1995). (B-C) Boxplot showing the distributions of numbers of mapped read-pairs and genes detected for each stage and protocol. (D) Sample correlation matrix showing the Pearson correlation coefficient between all pairwise combinations of samples. (E-G) Principal Component Analysis. (E) Principal Component (PC) 2 plotted against PC1. (F) PC3 plotted against PC1. (G) 3D rendering of the first 3 PCs. (H) Representation of expression profiles contributing to the first 6 PCs, produced by calculating the median expression value (normalised counts) for each sample for the 100 genes that contribute most to that PC.

**Figure 1-1.**
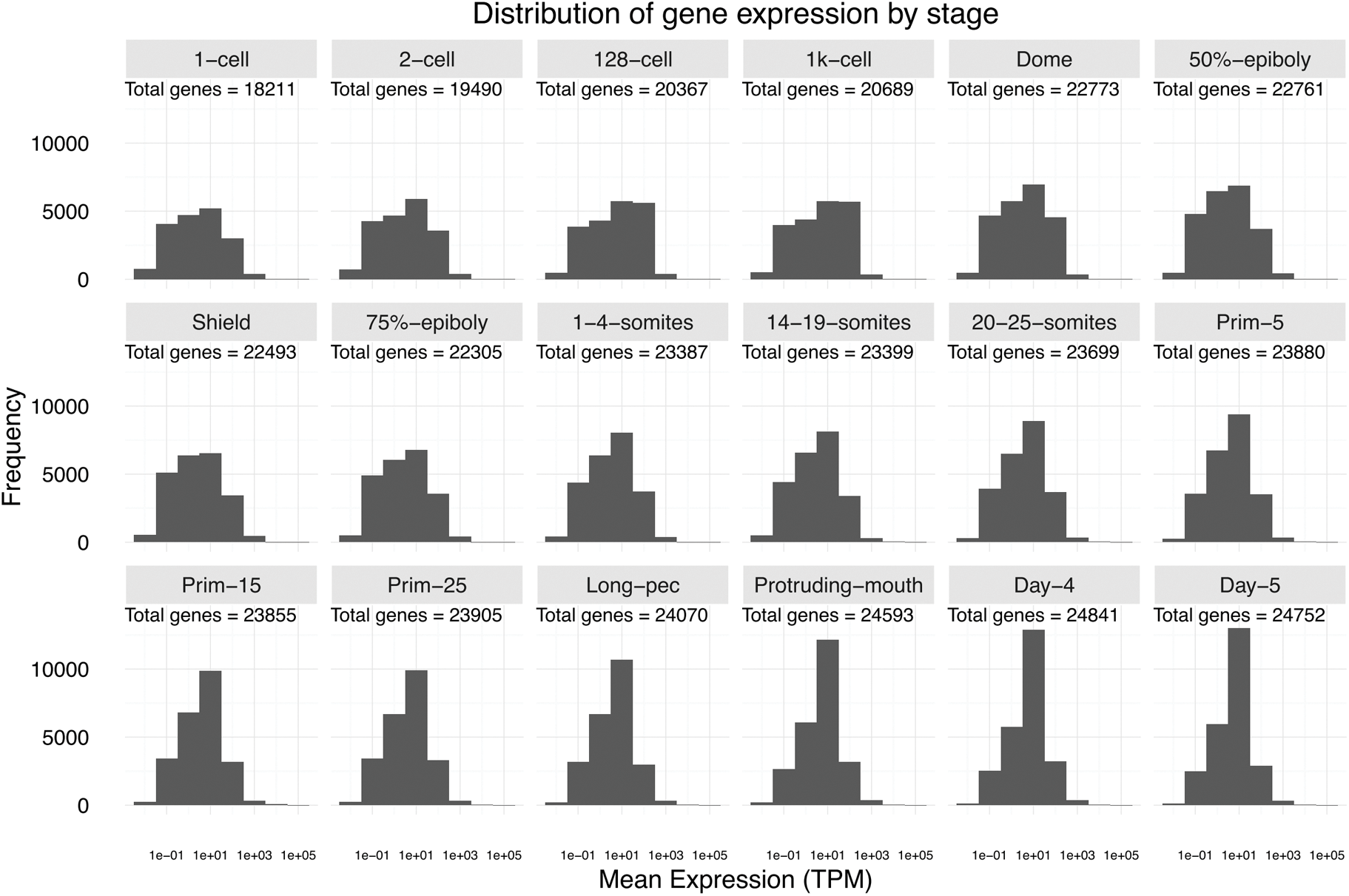
Distribution of gene expression levels in RNA-seq data by stage. Histograms (one per stage) showing the distribution of the mean expression level for each gene over the 5 samples within the stage. The total number of genes represented for the stage is shown on each panel.

**Figure 1-2.**
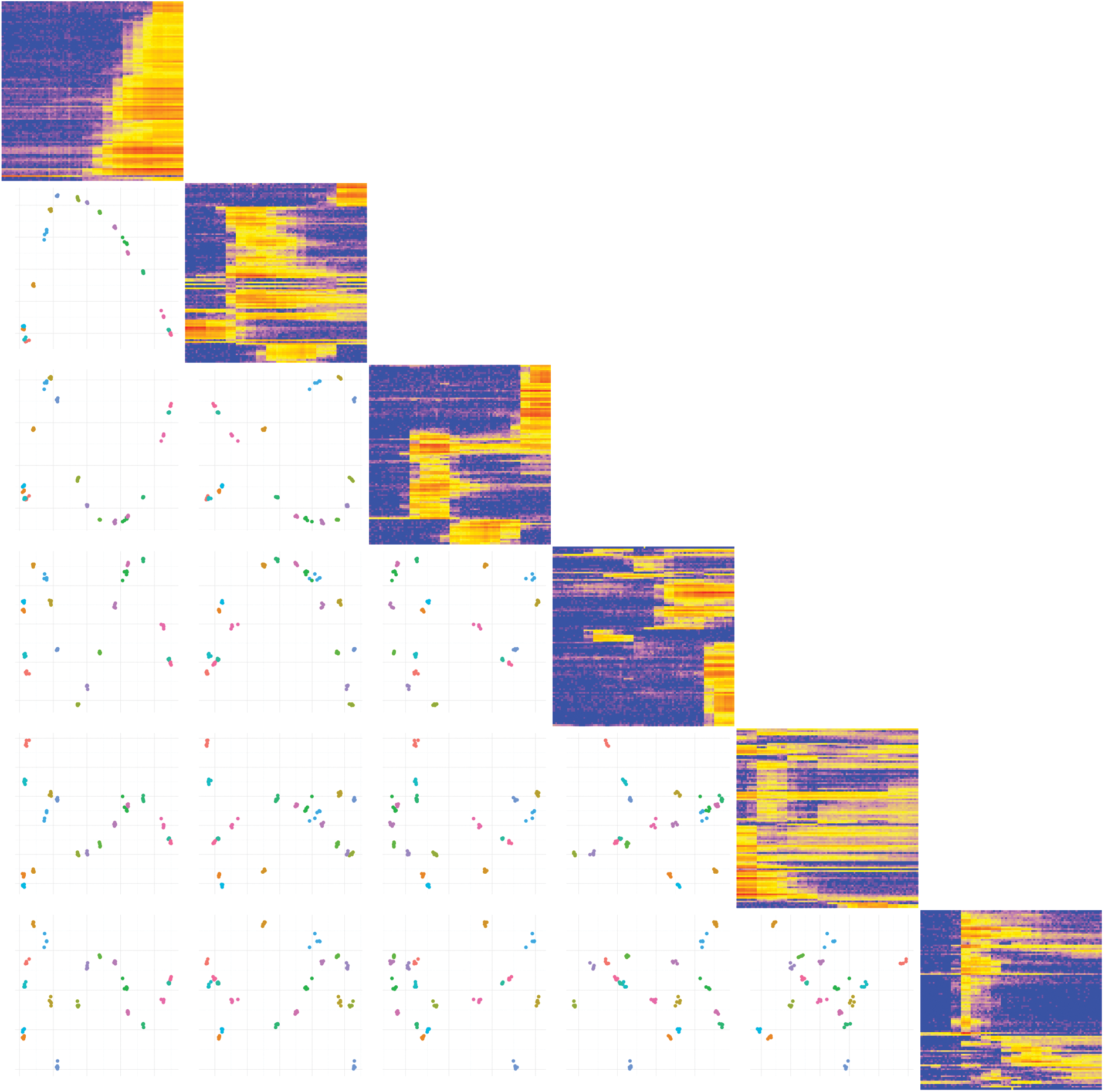
Matrix PCA plot. Plots showing the first six principal components (PCs) plotted against each other. The components are arranged in a matrix from PC1 (top and left) to PC6 (bottom and right). The plots on the diagonal show the expression profiles of the 100 genes contributing the most to that component. The plots below the diagonal show the appropriate PCs plotted against each other. For example, the plot in the third row and first column is PC1 (x-axis) plotted against PC3 (y-axis).

**Figure 1-3.**
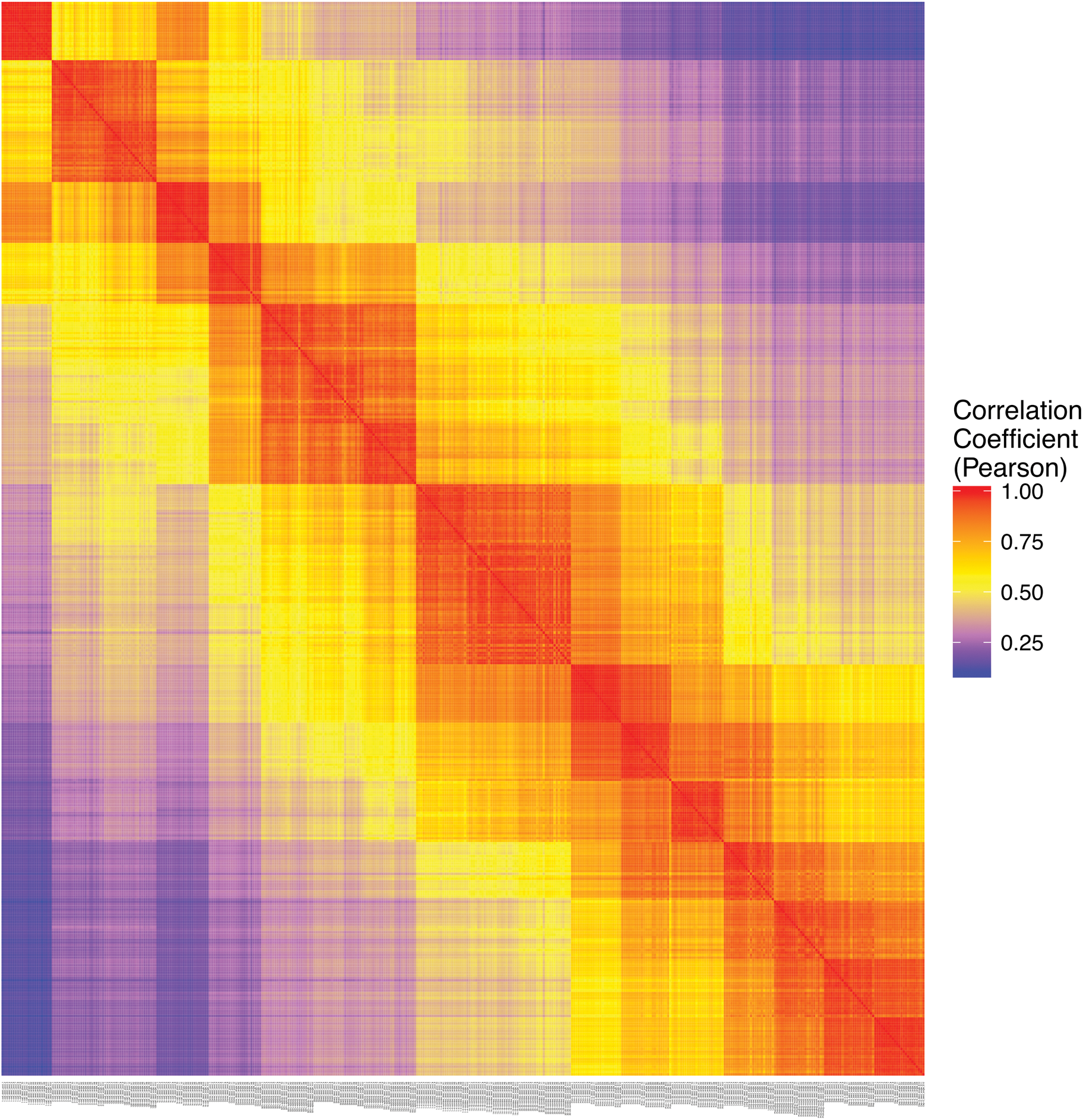
DeTCT sample Correlation matrix. Sample correlation matrix showing the Pearson correlation coefficient between each pair-wise comparison of samples.

**Figure 1-4.**
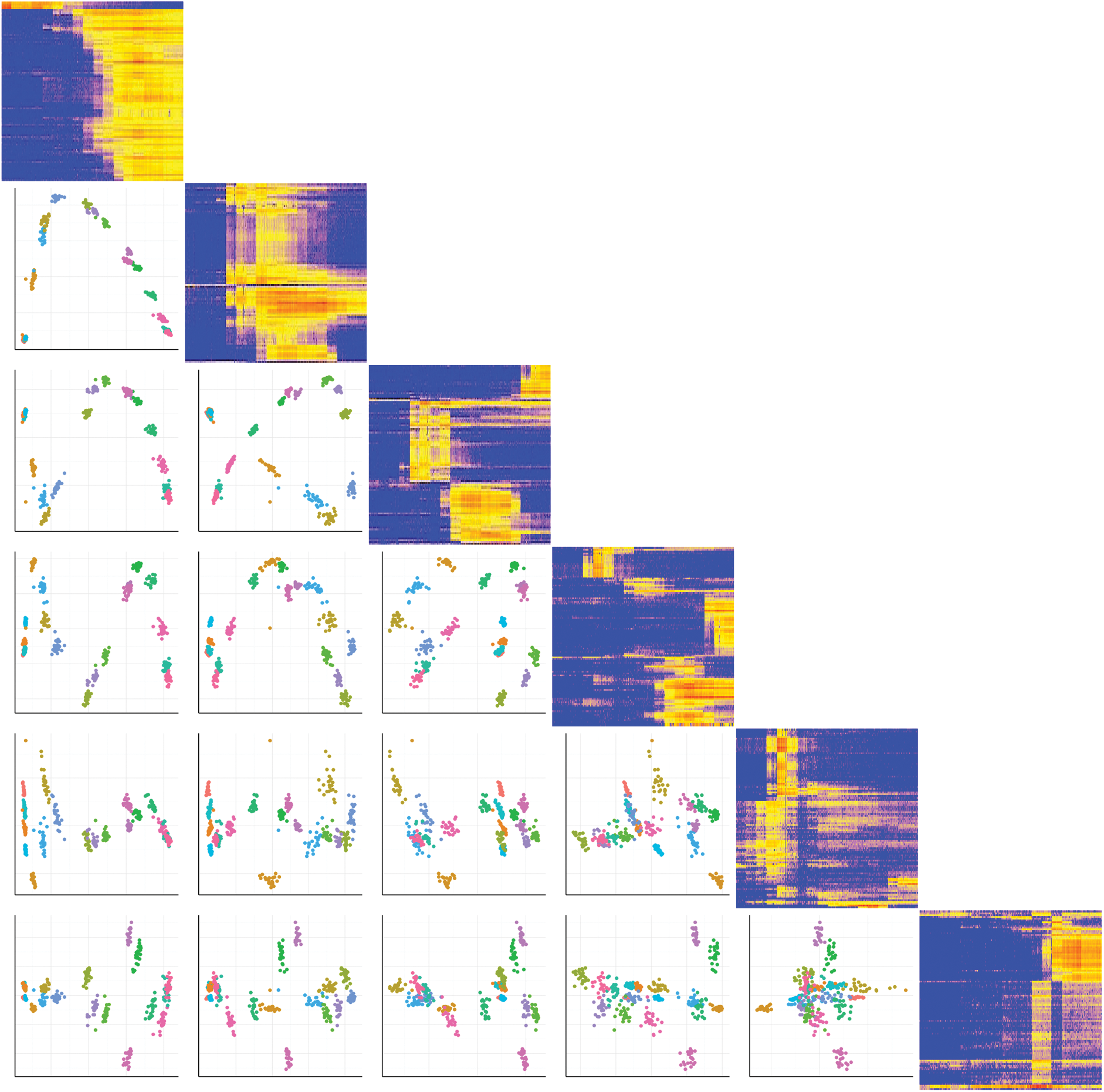
DeTCT Matrix PCA plot. Plot showing the PCA of the DeTCT data arranged in the same way as Figure 1-2.

**Figure 2.**
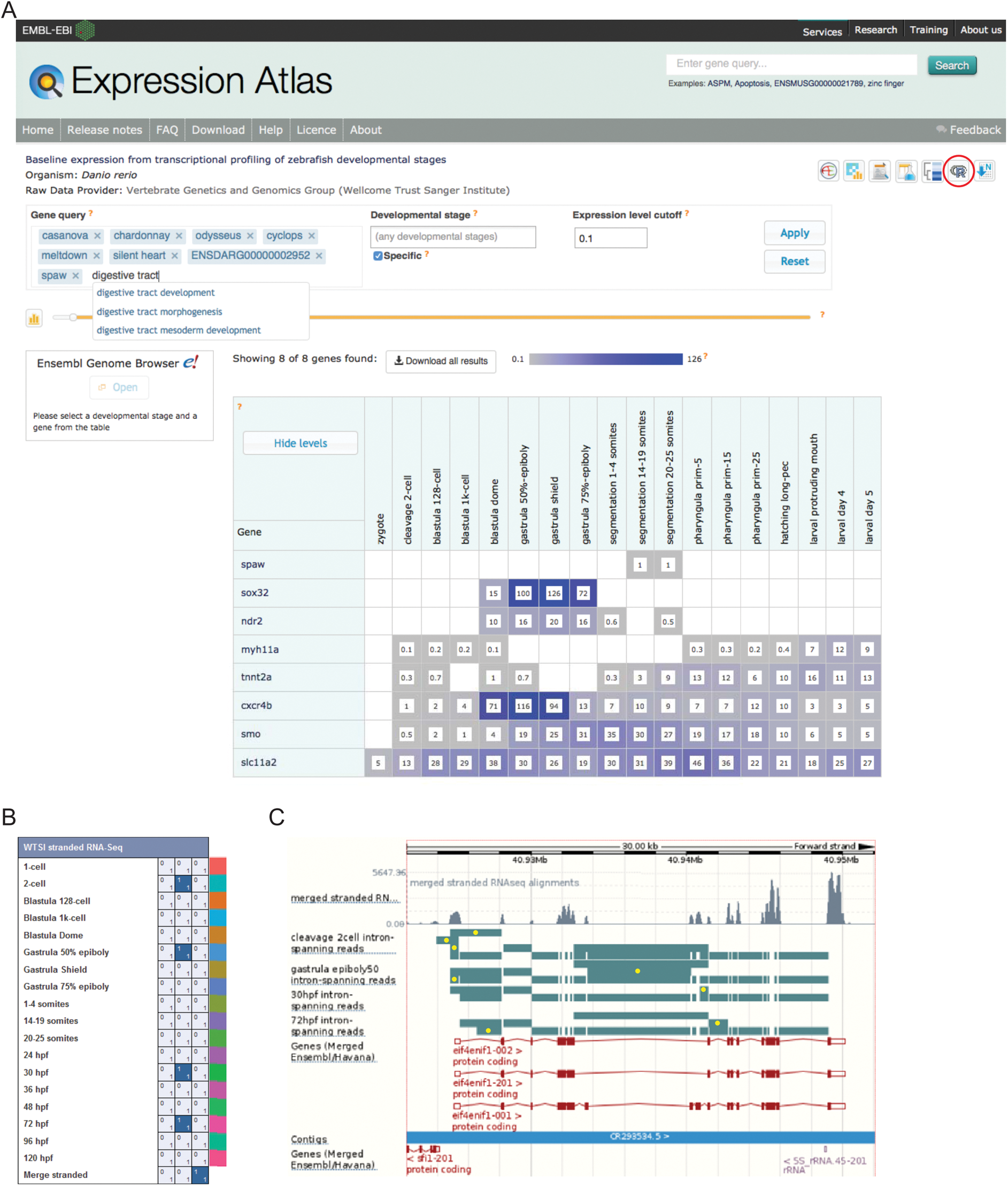
Data Access. (A) A screenshot from Expression Atlas release Dec 2016. Relative gene expression levels across all 18 stages are shown in FPKM. Genes can be searched for using Ensembl IDs, ZFIN gene IDs, gene/allele names and Gene Ontology terms. The data for the selected genes can be downloaded (red circle) as can the entire dataset. (B) A screenshot of Ensembl v87 showing the RNA-seq matrix with four intron tracks plus the merged RNA-seq data from exon read coverage from all 18 stages chosen for display. (C) A screenshot of Ensembl v87 showing the region surrounding the gene *eif4enif1* (chr 6:40.922-40.952 Mb). Exon read coverage from the merged 18 stages is shown as grey histograms. Four of the 18 intron-defining tracks from the RNA-seq matrix are shown. Introns are coloured in teal. Yellow dots indicate introns identifying a splice variant not shown by any of the three gene models shown in red. Note only the forward strand is shown in full and a gene model overlaps on the reverse strand.

### Temporal Map of Development

To visualise the expression of all genes across development we ordered the genes in our dataset by their stage of maximum expression (Figure 3A). This organises the genes temporally, broadly by expression pattern. These 18 groups of genes were then analysed for enrichment of Zebrafish Anatomy Ontology (ZFA) and Gene Ontology (GO) terms.

#### ZFA enrichment

The Zebrafish Anatomy Ontology (ZFA, Van Slyke et al., 2014) is a controlled vocabulary for describing the anatomy and development of zebrafish and is used by the zebrafish model organism database (ZFIN Howe et al., 2013a) to annotate the spatial expression patterns of genes from the literature and large-scale whole mount mRNA *in situ* hybridisation experiments (Rauch et al., 2003; Thisse et al., 2001). The results show enrichments for cell types and tissues that are appropriate to the stages of maximum expression (Figure 3A). There are no significant enrichments of terms during cleavage stages. At early gastrula stages terms such as hypoblast, organiser and hatching gland are enriched with endoderm, yolk and epidermis being enriched at late gastrula. During segmentation stages terms such as muscle, notochord and haematopoietic system are enriched and then at pharyngula and larval stages terms associated with the nervous system and the neural crest become enriched. By contrast, terms for mesodermal and endodermal organs such as kidney, spleen and liver are enriched at later stages (days 4 and 5). This demonstrates that these groupings of genes are consistent with what is already known about zebrafish development, thus validating our dataset.

#### GO enrichment

Enrichment analysis using the Gene Ontology (GO) also shows enrichments of terms in a stage-appropriate manner. At cleavage stages, terms linked to cell division, DNA repair, protein turnover and chromatin modification are enriched, consistent with the fast rate of cell division as well as the large changes in chromatin structure that occur at early stages. Subsequently, terms associated with determination of the embryonic axes and specification of the major tissue divisions are enriched. For example, genes linked to anterior-posterior, dorsal-ventral and left-right patterning are enriched from dome until prim-25 as well as genes annotated to terms such as gastrulation, heart development, muscle cell differentiation and brain development. At later stages, more specific terms such as synaptic transmission, potassium ion transport and glutamate receptor signalling are enriched. Coordinated expression of blood clotting factor genes leads to the appearance of blood coagulation as an enriched term at day 4.

Of the 23,642 genes 5,024 are uncharacterised (i.e. genes with automatically generated names such as zgc:* and si:ch*). Interestingly, at dome and gastrula stages, the proportion of these unnamed genes is much higher (29.2-39.4%) than at other stages (14.4-26.5%; Figure 3A). This could indicate a lack of orthologues and therefore that zebrafish gastrulation is enriched for the expression of genes exclusive to this species. However, we looked at the proportions of unnamed genes with orthologues in various species. Genes with no orthologues in any of the species were categorised as Danio, those with orthologues present in the fish species examined, but not the mammals were termed Teleost and those with orthologues present in the mammals as well were termed Vertebrate (although in many cases the identified last common ancestor is more basal than vertebrates e.g. Bilateria). The proportions of these categories are broadly constant across all 18 stages and there is not an increase of the Danio-specific genes at gastrulation (Supplemental Figure 3-1). This suggests that there is a large number of genes involved in vertebrate gastrulation that are yet to be characterised. The complete list of clusters along with the ZFA/GO term enrichment are provided in Supplemental Data Files 3-5.

**Figure 3.**
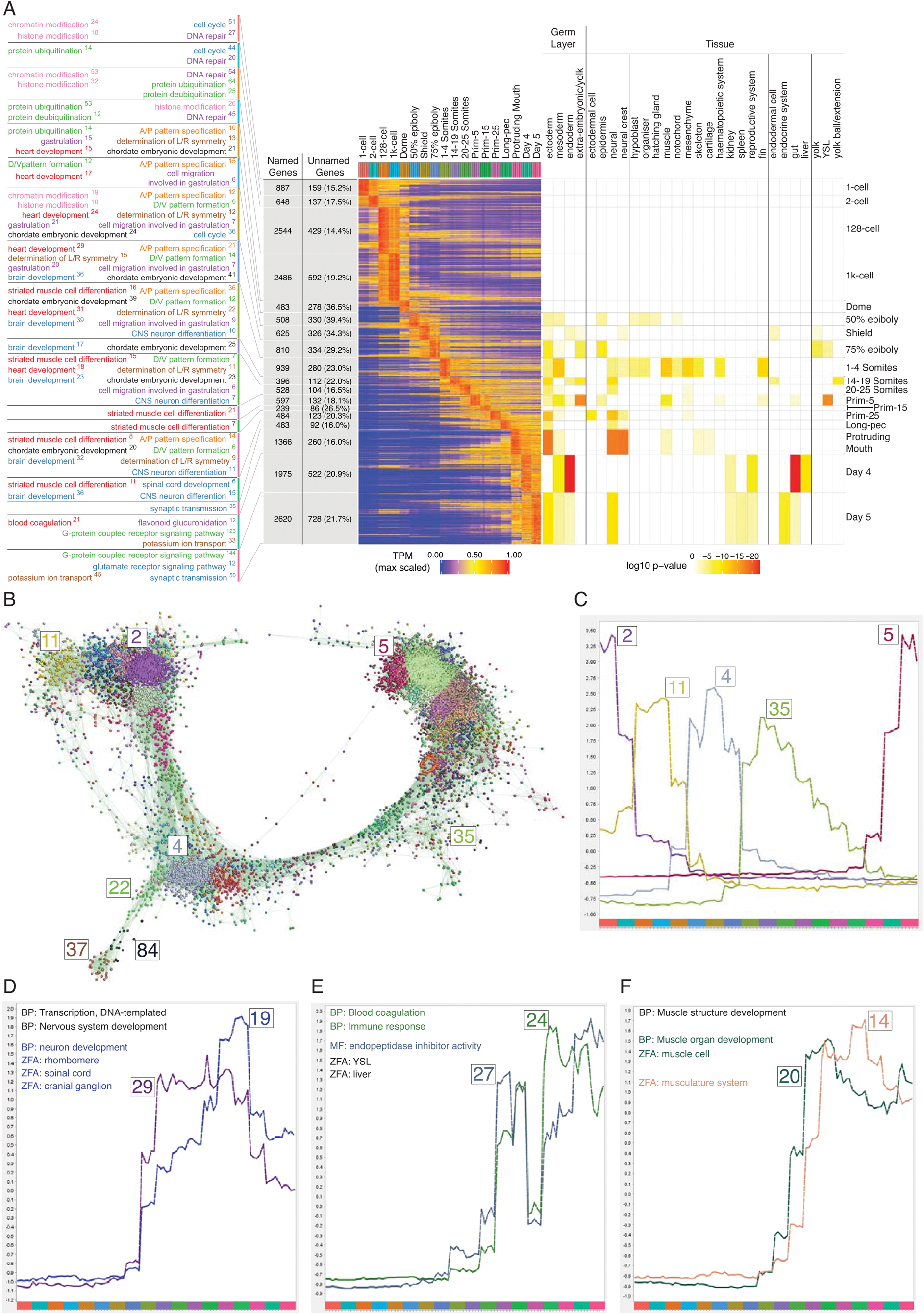
Clustering of expression patterns. (A) Heatmap showing the expression profiles of all the genes with appreciable expression in at least one stage. The expression values (Transcripts per Million, TPM) are scaled to the maximum expression for that gene across the time course and genes are organised by which stage the maximum expression occurs at. To the right is a heatmap of ZFA enrichments based on the genes assigned to each stage. To the left are a selection of the GO enrichments. The numbers of genes assigned to each GO term are shown as superscripts. Also shown are the numbers of named and unnamed genes assigned to each stage. (B) Network diagram showing the clustering produced by MCL on a network graph generated by linking genes that have at least one sample with 6 or more normalised counts, a Pearson correlation coefficient of > 0.94 and removing clusters with five genes or fewer. The numbers show the positions in the network diagram of the clusters shown in C and the ZnF clusters shown in Figure 4. (C-F) Example expression profiles displayed as lines showing the mean of mean-centred and variance-scaled normalised counts. (C) Example clusters showing the progression of expression peaks through development. (D-F) Expression profiles of example clusters with interesting GO/ZFA enrichments. Terms in black apply to all clusters in the diagram whereas terms specific to one cluster are shown in that colour.

**Figure 3-1.**
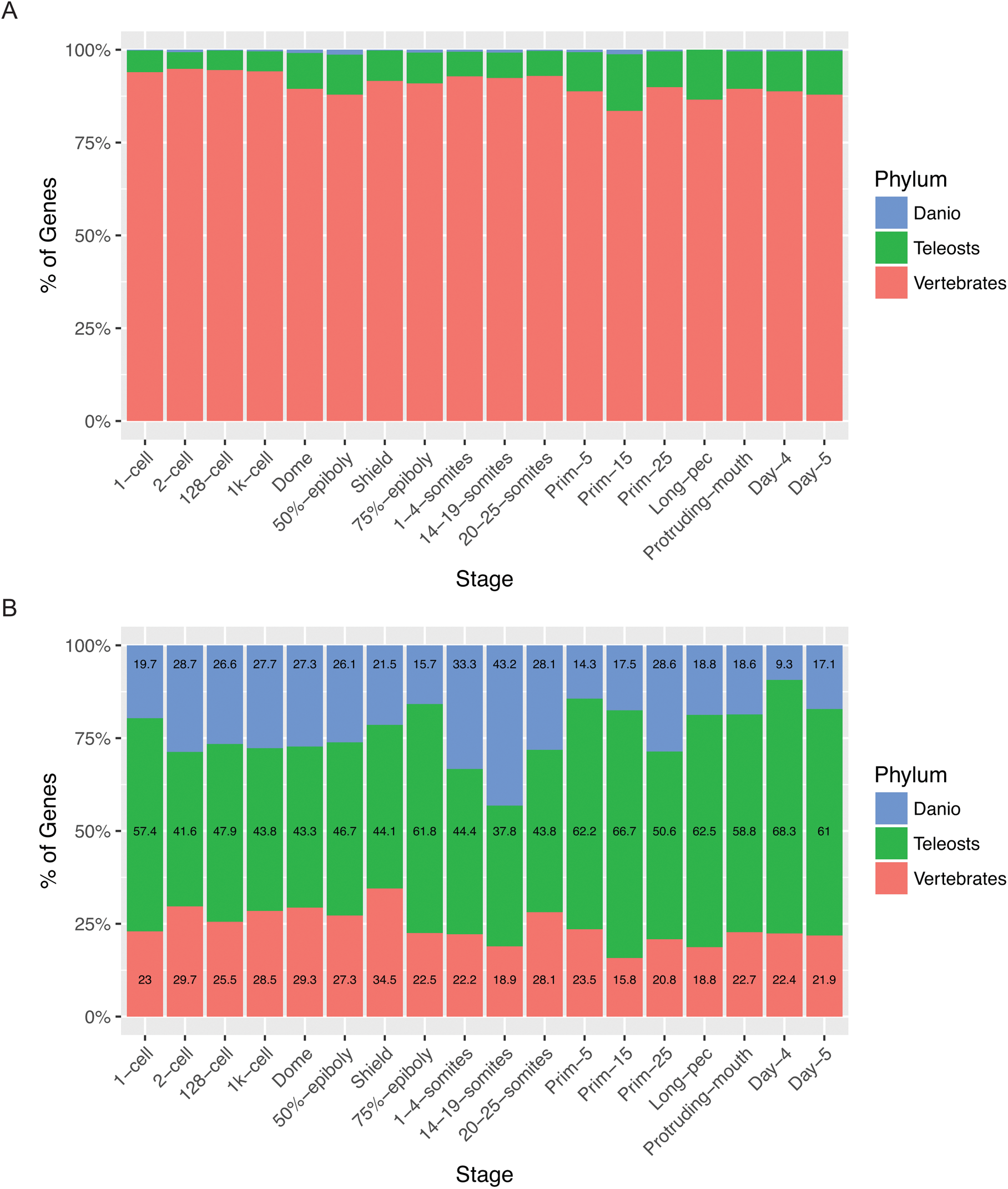
Relationship of named and unnamed genes to other species. Stacked bar charts showing the proportions of genes with orthologues at different taxonomic levels for Named (A) and Unnamed (B) genes. Orthologues for genes were obtained from the Ensembl Compara database and genes were then categorised based on which species they have orthologues in. Genes with no identifiable orthologue in any of the species examined were classed as *Danio* specific (top bar, blue). Those with orthologues in the six fish species (*Astyanax mexicanus*, *Gasterosteus aculeatus*, *Oryzias latipes*, *Takifugu rubripes*, *Tetraodon nigroviridis*, *Gadus morhua*), but not the mammals examined were labelled Teleosts (middle bar, green). Those with orthologues also present in the mammals surveyed (*Mus musculus*, *Rattus norvegicus* and *Homo sapiens*) were labelled Vertebrates (bottom bar, red).

### Graph-based clustering produces compact clusters of highly biologically related genes

While the maximum expression stage groupings provide a high-level, temporally organised view of development, smaller more specific groupings would provide more insight especially into the role of uncharacterised genes. To get smaller clusters that are more likely to be highly biologically related we used the BioLayout *Express*^3D^ software (http://www.biolayout.org/) (Theocharidis et al., 2009) to cluster and visualise the dataset (Figure 3B). The software first builds a network graph in which genes are nodes with edges between those whose expression is correlated above a given threshold. It then uses the Markov Cluster Algorithm (MCL, Enright et al., 2002; van Dongen, 2000) to find strongly connected components within the graph. These are clusters of genes that are more highly connected with each other than with the rest of the graph. This produces 252 clusters, the vast majority of which (238) have fewer than 100 genes (Figure 3B). Broadly speaking, the clusters on the left-hand side of the graph represent genes expressed at early stages, whereas towards the right of the graph the expression profiles are progressively later in development (Figure 3B-C). For example, cluster 2 contains genes that are maternally supplied and then degraded, cluster 11 has genes that accumulate after the 2-cell stage (possibly by control of polyadenylation) and are then cleared. To test whether this is an artefact of the normalisation process, we also normalised the counts using the ERCC spike-in RNAs included in every sample. The expression profiles are similar, although the ERCC normalisation introduces more variability from embryo to embryo within the same stage (Figure 3-2). After the maternal stages, cluster 4 genes are expressed at the time of zygotic genome activation, cluster 35 genes are expressed during segmentation stages and genes from cluster 5 are not expressed until the last time points in the study (Figure 3C).

**Figure 3-2.**
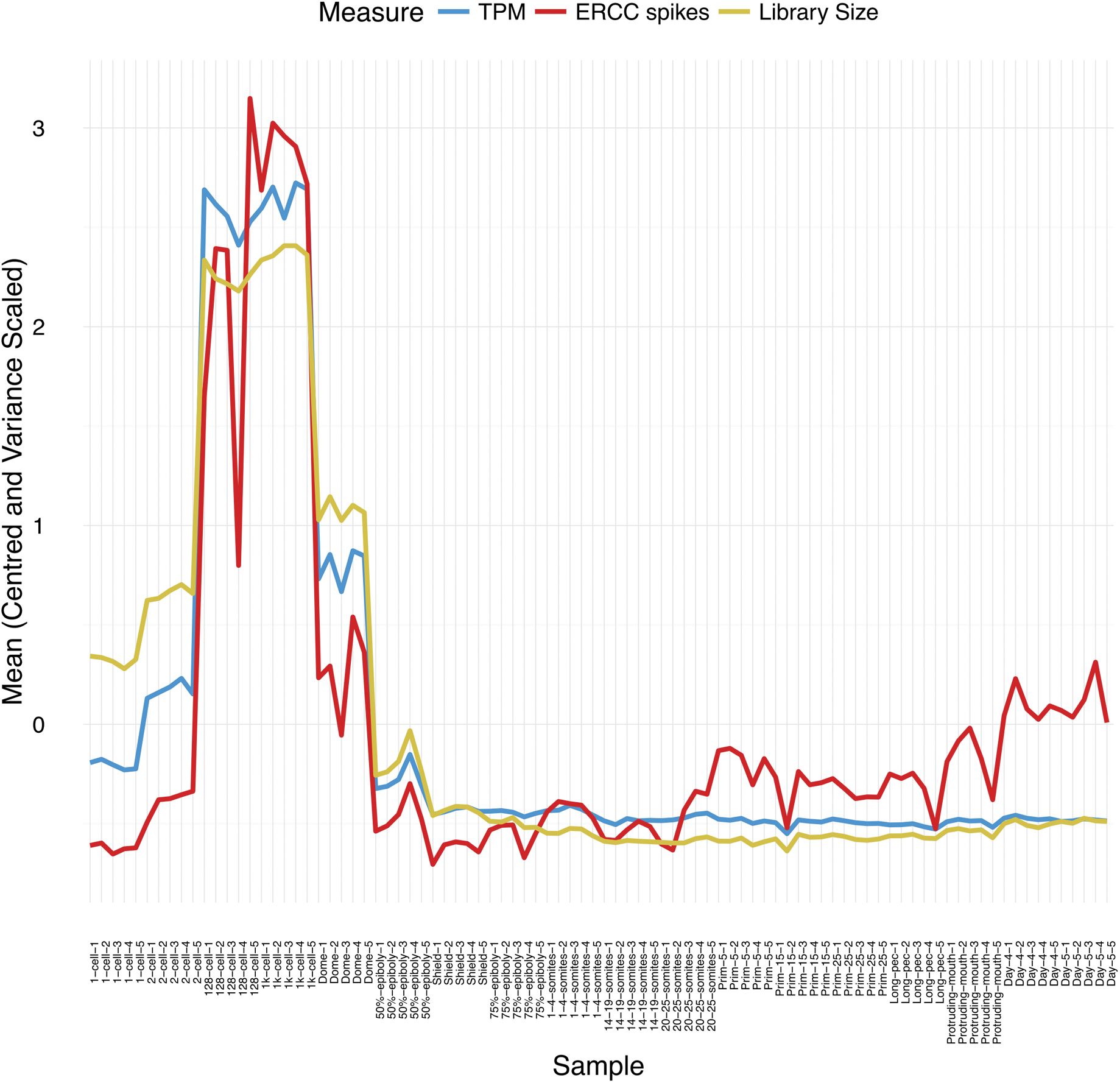
Effect of normalisation on expression profile of cluster 11 genes. Counts were normalised by DESeq2 using either the ERCC spike-in RNAs (ERCC spikes, red line) or all the counts for each sample (Library size, yellow line). As a comparison, TPM is also shown (blue line).

We again used GO and ZFA enrichment to analyse the composition of the genes within the clusters. For example, cluster 19 contains large numbers of transcription factors from the *dlx*, *hox*, *irx* and *pax* gene families that are expressed from 1-4 somites onwards and is associated with GO terms for transcription and nervous system development and ZFA terms such as rhombomere, spinal cord and cranial ganglion (Figure 3D). Cluster 29, with similar GO enrichments, has an overlapping expression profile but an earlier and broader peak. It contains genes such as *nkx2.1*, *gli3*, *sox1b* and *hoxb* genes. Clusters 24 and 27 show a sharp dip in expression at 36 hpf and contain complement, clotting factor genes and serpins. Both clusters have significant enrichments for the terms blood coagulation and endopeptidase inhibitor activity (Figure 3E). (Figure 3F). The information on cluster assignments and GO/ZFA enrichments are provided in Supplemental Data Files 6-9.

### A burst of transcription of highly related zinc finger proteins during zygotic genome activation

During examination of the clusters produced by BioLayout, we noticed several clusters that appeared to be enriched for a family of zinc finger (ZnF) proteins located on chromosome 4 (Howe et al., 2016). To test systematically if any chromosomes were over-represented in any of the clusters, we performed a binomial test for each chromosome for each cluster that contained more than five genes from that chromosome. Table 1 shows the clusters and chromosomes that are significantly enriched by more than twofold. This identifies four clusters (Figure 3B, clusters 4, 22, 37 and 84) with more chromosome 4 genes than would be expected by chance given the size of the clusters. Most of the chromosome 4 genes within these clusters are predicted to encode proteins with ZnF domains (Supplemental Data File 10).

The expression profiles of these ZnF genes show a burst of expression from dome to 75% epiboly (4.3-8 hpf) and several of them show appreciable RNA levels at the 128- and 1000-cell stages (Figure 4A-B, Supplemental Figure 4-1). Cluster 37 shows a particularly sharp peak with a very large increase from the 1000-cell stage to dome and an almost equal decrease from dome to 50% epiboly. This suggests that these ZnF genes are co-ordinately expressed at the onset of zygotic transcription. To investigate whether this is a global effect on all the genes on the long arm of chromosome 4 we looked at the expression profiles of NLR family genes, another large family of proteins present on chromosome 4. These genes are present in the same region of chromosome 4 as the ZnF genes although their distribution is slightly different, with more NLR genes towards the end of the chromosome (Figure 4F-G, Supplemental Figure 4-2). Of the 312 NLR genes present in the Ensembl version 85 genebuild, 149 have appreciable expression during this time course and they are not expressed in the same pattern as the ZnF genes (Figure 4C).

**Figure 4.**
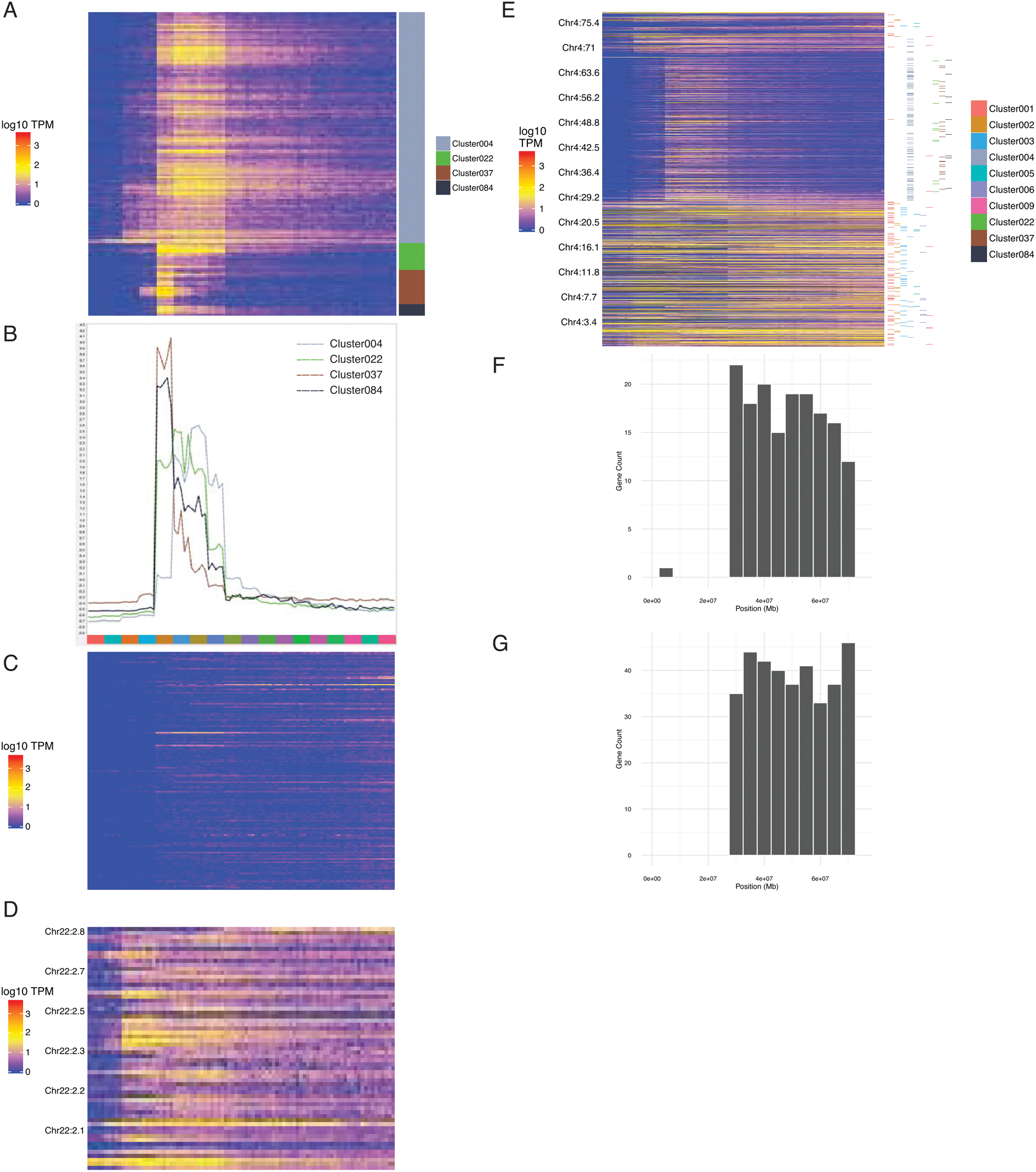
ZnF genes expressed at the onset of zygotic transcription. (A) Heatmap showing the expression profiles (log10 TPM) of the ZnF genes from clusters 4, 22, 37 and 84. (B) Expression profiles of clusters 4, 22, 37 and 84 shown as average expression (mean centered and variance scaled). (C) Heatmap showing the expression profiles (log10 TPM) of the NLR genes present in GRCz10. (D) Heatmap showing the expression of a cluster of ZnF genes on chromosome 22 displayed in chromosome order. (E) Heatmap showing the expression of all genes on chromosome 4 (log10 TPM) in chromosomal order with cluster assignments shown on the right. (F-G) Histograms showing the distribution of ZnF (F) and NLR (G) genes on chromosome 4.

**Figure 4-1.**
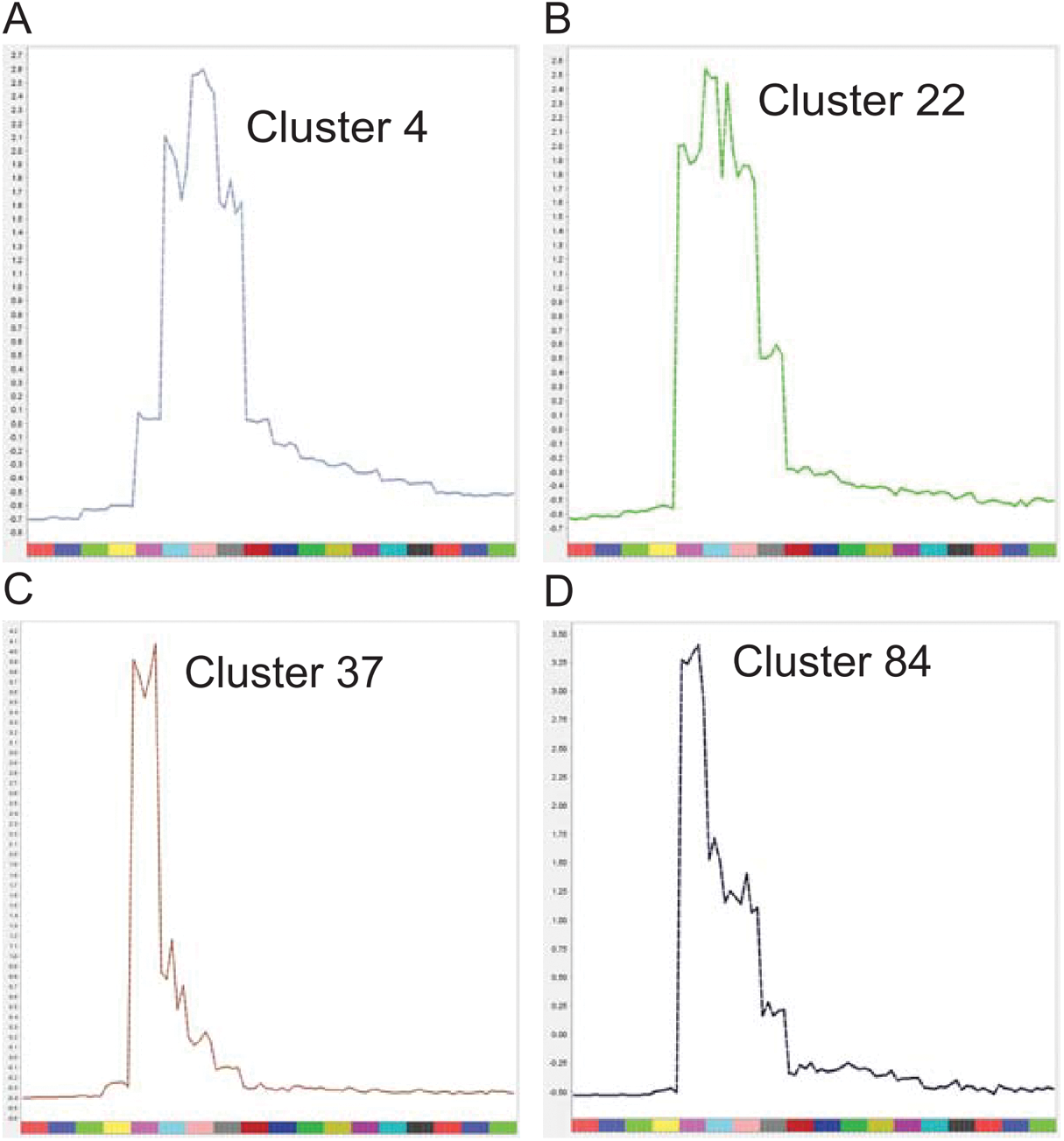
BioLayout cluster profiles. Individual expression profiles of clusters 4 (A), 22 (B), 37 (C) and 84 (D) shown as average expression (mean centered and variance scaled).

**Figure 4-2.**
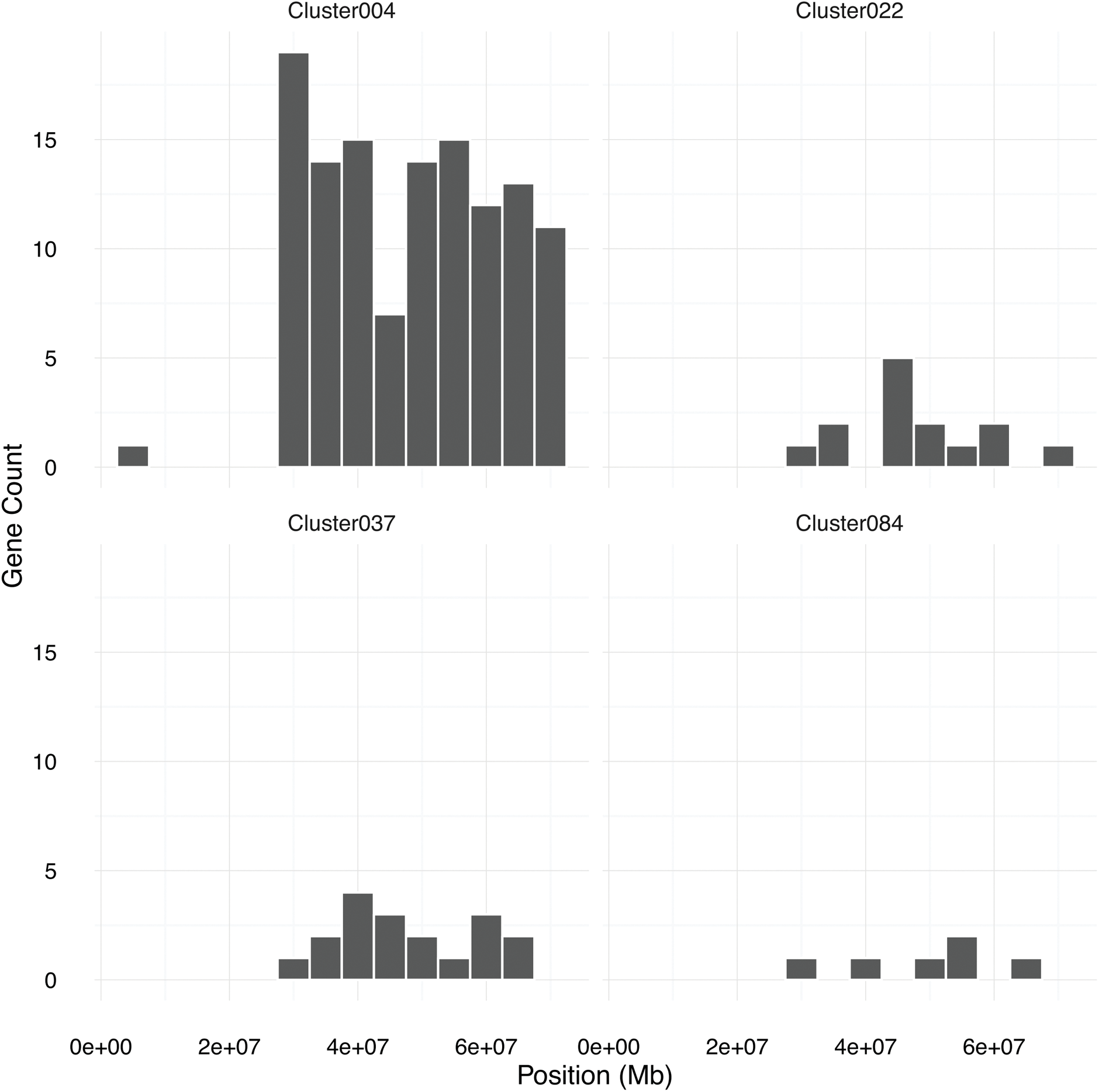
Chromosomal distribution of the chromosome 4 ZnF genes. Individual histograms split by cluster showing the chromosomal distribution of the ZnF genes from clusters 4, 22, 37 and 84.

This effect can even be seen on the chromosomal scale. A plot of the expression profiles of all the genes across chromosome 4 in chromosomal order shows a domain on the long arm that is enriched for these ZnF genes (Figure 4E) that share a similar expression pattern.

**Table 1.** BioLayout clusters with significant chromosomal enrichments (adjusted p-value < 0.05 and log2 fold enrichment [log2FE] > 1).

| Cluster | Chromosome | Count | Expected | log2FE | Adjusted pvalue |
| --- | --- | --- | --- | --- | --- |
| Cluster023 | Chr1 | 9 | 2.25 | 2.00 | 1.161e-02 |
| Cluster004 | Chr4 | 129 | 47.29 | 1.45 | 7.434e-24 |
| Cluster022 | Chr4 | 14 | 5.03 | 1.48 | 1.156e-02 |
| Cluster037 | Chr4 | 23 | 3.35 | 2.78 | 9.033e-14 |
| Cluster084 | Chr4 | 9 | 1.48 | 2.61 | 1.659e-04 |
| Cluster074 | Chr12 | 8 | 0.48 | 4.05 | 6.337e-07 |

#### Chromosomal expression domains

Since the expression pattern of the chromosome 4 ZnF genes is obvious on a chromosome level plot, we wondered if it was possible to find any other chromosomal regions where the genes share a common expression pattern. To investigate this, we plotted the expression of all the genes for each chromosome in chromosomal order (Supplemental Figure 4-3A). This led to the identification of another region with genes containing zinc finger domains, this time on chromosome 22 (Figure 4D). These genes begin to be expressed at a similar stage to the chromosome 4 ZnF genes, although they tend to be more widely expressed over the time course (Figure 4D).

There are also several other domains of co-expressed genes that can be clearly seen at the chromosome level, such as the gamma crystallin family of genes on chromosome 9, that are all highly expressed at the end of the time series (Supplemental Figure 4-3B).

Interestingly, there is a small region on chromosome 12 where the genes are all expressed from late gastrula through to the end of the pharyngula period (Supplemental Figure 4-3C). The genes in this region all have clone-based names apart from *ctslb* (ENSDARG00000039173, also known as *hatching gland 1*, *hgg1*). All of them appear in the same BioLayout cluster (cluster 74), which has an overrepresentation of genes on chromosome 12 (Table 1). This appears to be a fish-specific expansion of cathepsin L. Similar expansions have happened in multiple different lineages including rodents (http://www.ensembl.org/Mus_musculus/Gene/SpeciesTree?db=core;g=ENSMUSG00000021477;r=13:64363214-64370306;t=ENSMUST00000021933).

On chromosome 11 there is a cluster containing hairy and enhancer of split-related genes and a set of unnamed genes all expressed from late gastrula through segmentation and pharyngula stages (Supplemental Figure 4-3D). Finally, two clusters of genes containing histone domains can be seen on chromosomes 7 and 25 (Supplemental Figure 4-3E-G).

**Figure 4-3.**
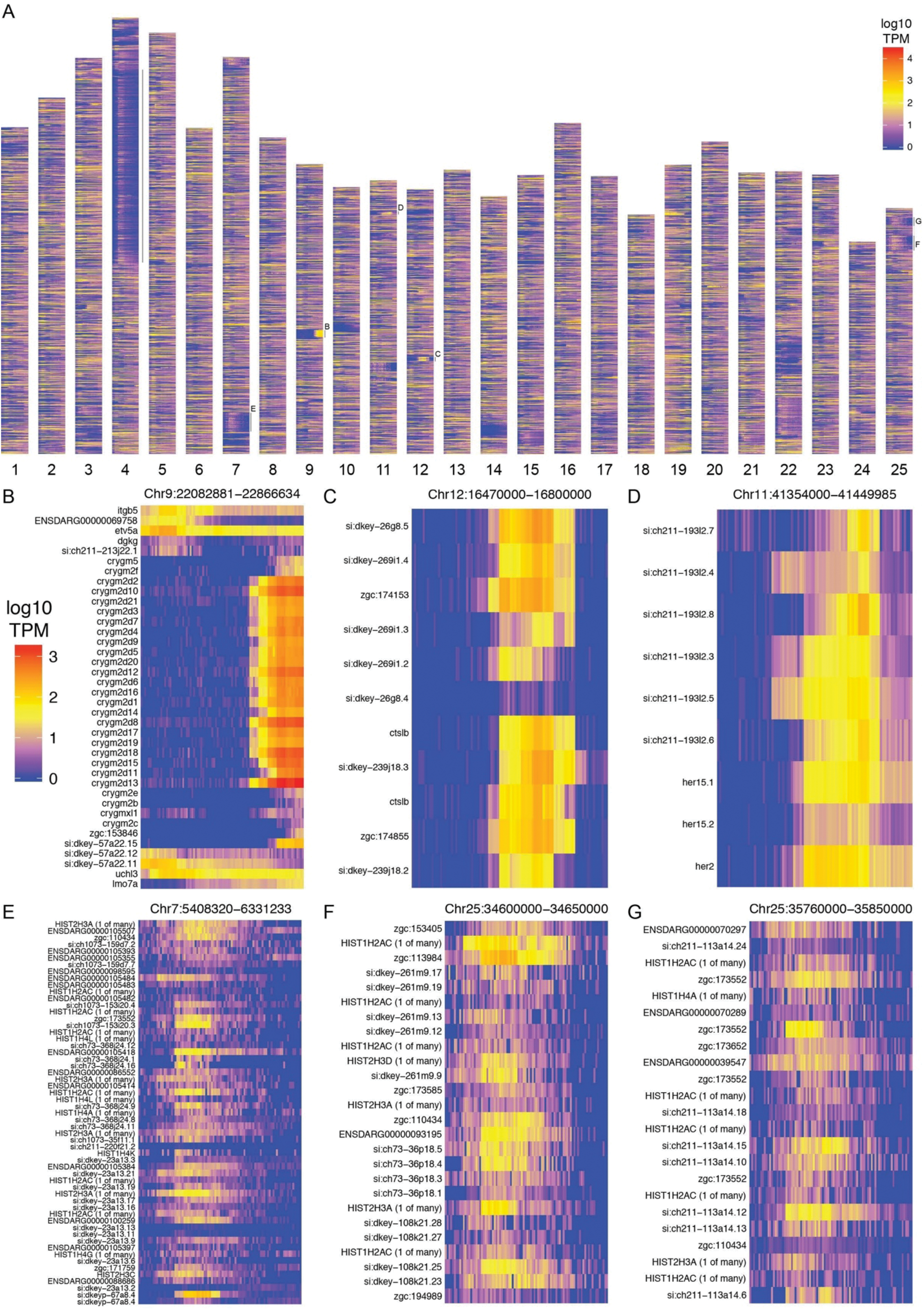
Chromosomal expression domains. (A) Heatmaps showing the expression profiles (log10 TPM) of genes in chromosomal order. The unlabelled line next to chromosome 4 marks the location of the ZnF genes shown in Figure 4. (B-G) Expression profile heatmaps of the regions indicated by letters in (A).

### Conservation/Divergence of the expression profiles of paralogous genes

Teleost fish underwent a third round of whole genome duplication compared to the two rounds of tetrapods. During rediploidisation, most of the duplicate genes were lost, however a significant proportion were retained and are now present as paralogous pairs. Paralogous genes can have conserved expression patterns but often the expression patterns have diverged so that the paralogues function at different times or in different tissues. To investigate whether we can see this in our data, which has only stage and not tissue specificity information, we produced a list of one-to-one teleost-specific paralogues pairs using data from Ensembl (3144 gene pairs with expression in our data).

We calculated the Pearson correlation coefficients between the normalised counts for each paralogous pair as a measure of how similar the expression profiles are. A histogram of the correlation coefficients shows a distribution skewed to high positive correlations (Figure 5A) when compared to a random sample of gene pairs of similar size (Figure 5B). Despite this skew, there is a significant proportion of the distribution with low or negative correlation coefficients, suggesting that this may represent a set of genes whose expression patterns have diverged (50% of the distribution is below 0.45).

Examples of highly correlated and negatively correlated genes are shown in Figure 5C-G and Supplemental Figure 5-1. To get an estimate of how many of these gene pairs have divergent spatial expression patterns, we used the ZFA annotation provided by ZFIN. There are 683 gene pairs where there is anatomical expression information for both genes. For each pair we counted up the intersection of all the pairs of stage ID and anatomy term that make up the expression annotations. For the pairs studied, 83.7% (572 pairs) have an intersection of less than 50%.

**Figure 5.**
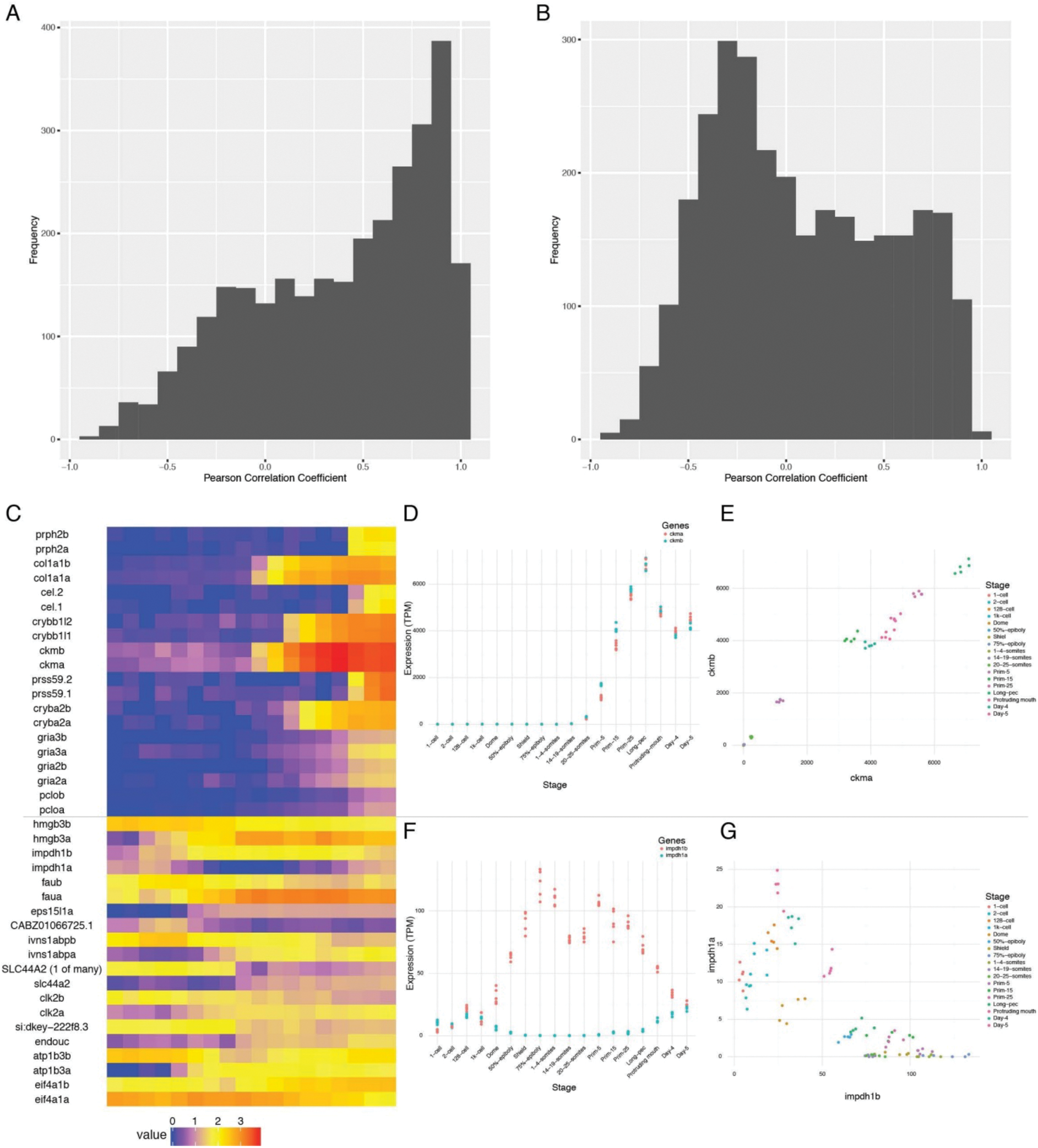
Conservation/divergence of paralogous genes. (A-B) Histograms of Pearson correlation coefficients among one-to-one paralogous gene pairs (A) and a random sample of genes (B). (C) Heatmap showing the expression of the gene pairs with the top 10 highest correlation coefficients (top half) and the top 10 most negative correlation coefficients (bottom half). (D-E) Plots showing the expression of *ckma/b* by stage (D) and plotted against each other (E). (F-G) Plots showing the expression of *impdh1a/b* by stage (F) and plotted against each other (G).

**Figure 5-1.**
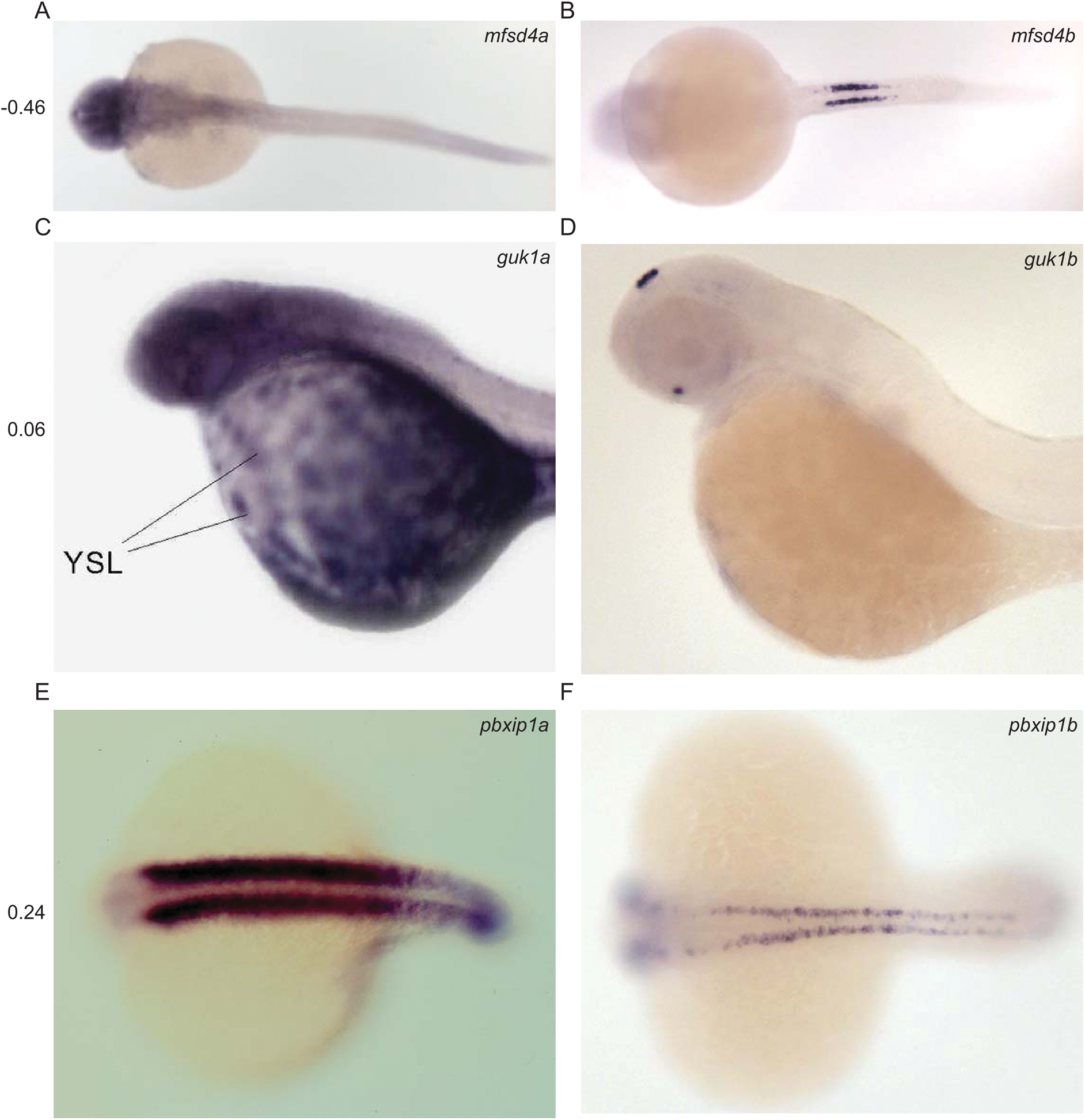
Divergent expression of paralogous pairs. Images from ZFIN (ZFIN.org) showing examples of paralogous gene pairs with diverged expression patterns. The Pearson correlation coefficient between the RNA-seq expression profiles of each pair is shown to the left. (A-B) *mfsd4a/b*. (C-D) *guk1a/b*. (E-F) *pbxip1a/b*.

### Alternative 3′ end use

To catalogue alternative 3′ end usage across development we produced 3′ pulldown data for the same time points as the RNA-seq. There are multiple possible consequences of alternative 3′ ends, such as different/extra 3′ UTR sequence (Figure 6A). This could provide different binding sites for RNA-binding proteins or regulation by microRNAs. For example, one transcript may contain a microRNA binding site lacking in another, changing post-transcriptional regulation. Other possibilities include different/extra coding sequence at the 3′ end of transcripts, leading to proteins that contain different domains and therefore different molecular functions.

Across all 423 samples over all the stages, there were 253,627 regions identified, where each region may be associated with multiple 3′ ends. However, many of these are false positives due to poly(T) pulldown from genomic poly(A) tracts within transcripts. These regions are then filtered by various criteria such as distance to an annotated gene and genomic context in an attempt to remove artefactual regions (Figure 6B and Methods), which reduces the set to 37,724 regions. For this analysis, we increased the stringency of filtering to reduce the inclusion of false-positive ends and produce a high-confidence set of 3′ ends to work from. This stricter set contains 8358 regions. Once the regions have been associated with annotated genes, 1551 genes have two or more 3′ ends.

This high-confidence set has allowed us to define 3′ ends of genes that are alternatively used throughout development. For example, *pdlim5b* (ENSDARG00000027600) has three alternative 3′ ends which are used at different times during development (Figure 6C-E). During cleavage and gastrula stages, one particular end is used (end 2 in Figure 6C), but all three are observed during segmentation stages. Ends 1 and 2 would be predicted to produce the same protein with different 3′ UTRs, allowing for differential post-transcriptional regulation.

However, end 3 is predicted to produce a protein which lacks the C-terminal zinc finger domain, allowing for this alternative transcript to have a completely different molecular function.

**Figure 6.**
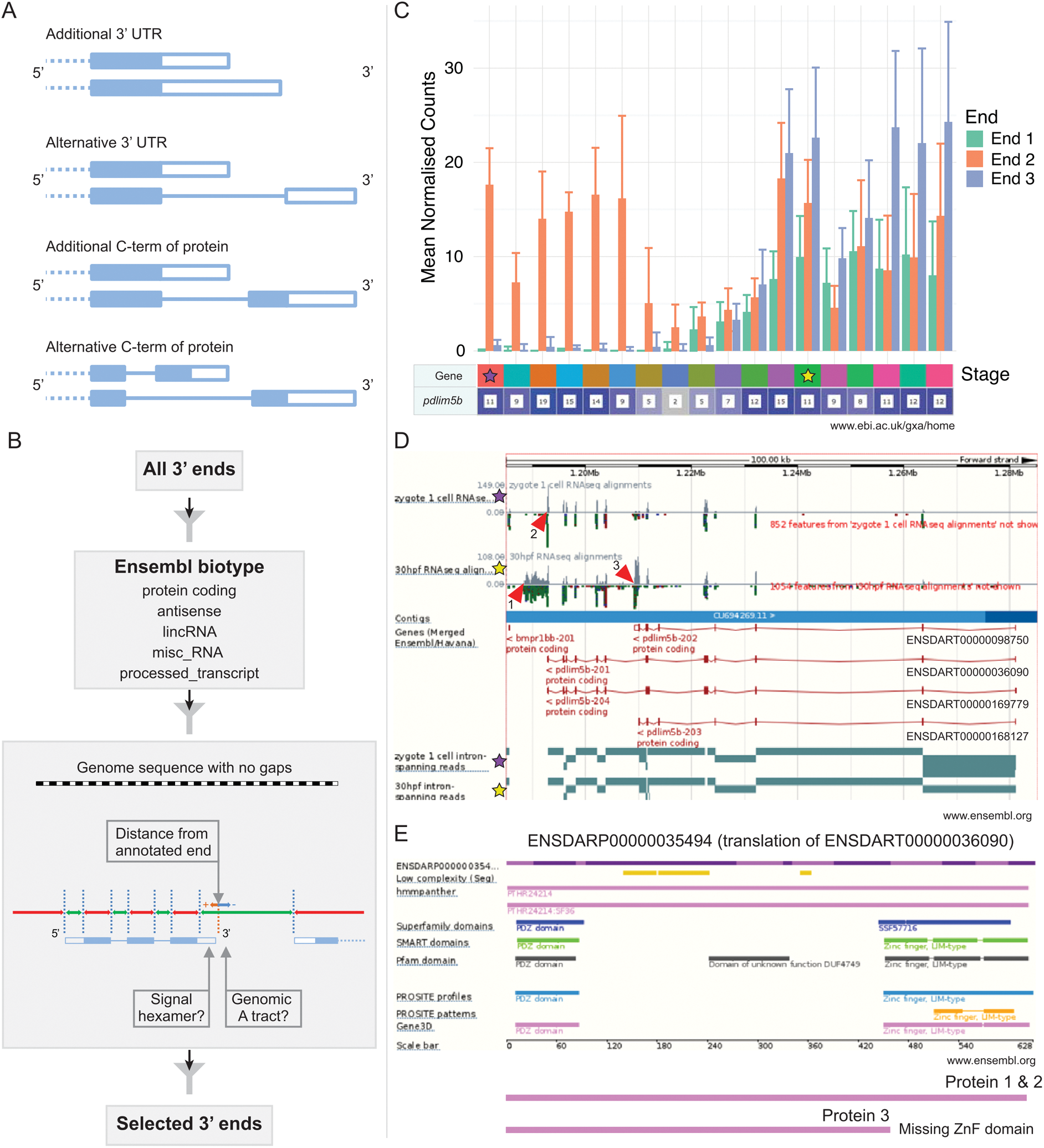
Alternative 3’ end use during development. (A) Diagrams showing the alternative possible outcomes of differential 3’ end usage. Only the 3’ end of the gene is shown where the exons are boxes, the introns are represented as lines and filled boxes are coding. The dotted line represents the remaining 5’ end of the gene. (B) Flowchart showing the DeTCT 3’ end filtering strategy. (C) Graph showing the mean normalised counts by stage for the three alternative 3’ ends (shown in D) of the *pdlim5b* gene. Error bars represent standard deviations. Underneath is a screenshot from Expression Atlas (Sept 2016) showing the overall FPKM for *pdlim5b*. The stages for which exon and intron data are shown in D are denoted with stars. (D) Screenshot from Ensembl (release 87) showing the *pdlim5b* gene (10:1185001-1285000). The purple stars mark exon read counts for the 1-cell stage at the top and intron spanning read counts at the bottom. Similarly, prim-15 counts are marked by yellow stars. Red arrowheads point to the genomic positions of the three transcript ends. Note 3’ UTR resulting in end 1 is not annotated in this gene build. Four transcripts are labelled on the right hand side. (E) A snapshot of Ensembl release 87 protein view showing *pdlim5b* transcript ENSDART00000036090.6 from N-terminus to C-terminus. The protein domains missing if the transcript finished at end 3 are shown below.

### Novel primary miRNA transcripts

Along with cataloguing 3′ ends, we have performed RNA-seq on mutants for two genes involved in microRNA (miRNA) maturation, in order to annotate the polyadenylated primary miRNA transcripts. Mature miRNAs are released from longer primary transcripts (pri-miRNAs) by a series of processing steps (Denli et al., 2004; Gregory et al., 2004; Grishok et al., 2001; Hutvagner et al., 2001). First, precursor hairpins (pre-miRNAs) are sliced from the host transcript in the nucleus by the microprocessor complex, consisting of the proteins Drosha (Rnasen) and Dgcr8 (Pasha). Subsequently the hairpin is cut by Dicer1 in the cytoplasm, freeing the miRNA duplex.

Although much effort has been dedicated to sequencing and annotating the mature miRNAs themselves, the annotation of many primary miRNA genes is likely hindered by their rapid nuclear processing. Reasoning that the depletion of *drosha* and *dgcr8* may enrich the transcriptome for unprocessed pri-miRNAs (Chang et al., 2015), we sequenced RNA derived from *dgcr8* and *drosha* wild-type, heterozygous and homozygous mutant embryos to determine whether this would allow us to improve the current pri-miRNA annotation. Following the assembly of the transcriptome, we were able to better define a large number of novel pri-miRNA loci not annotated in the current Ensembl release (v87) (Fig. 7). The v87 release includes 313 pri-miRNA transcripts overlapping 220 annotated miRNA loci. Our assembly contains 398 pri-miRNA transcripts which overlap 283 miRNA loci, 94 of which are not currently associated with a pri-miRNA (Figure 7D, Supplemental Data File 11 & 12). These will be incorporated into a future Ensembl gene set update.

**Figure 7.**
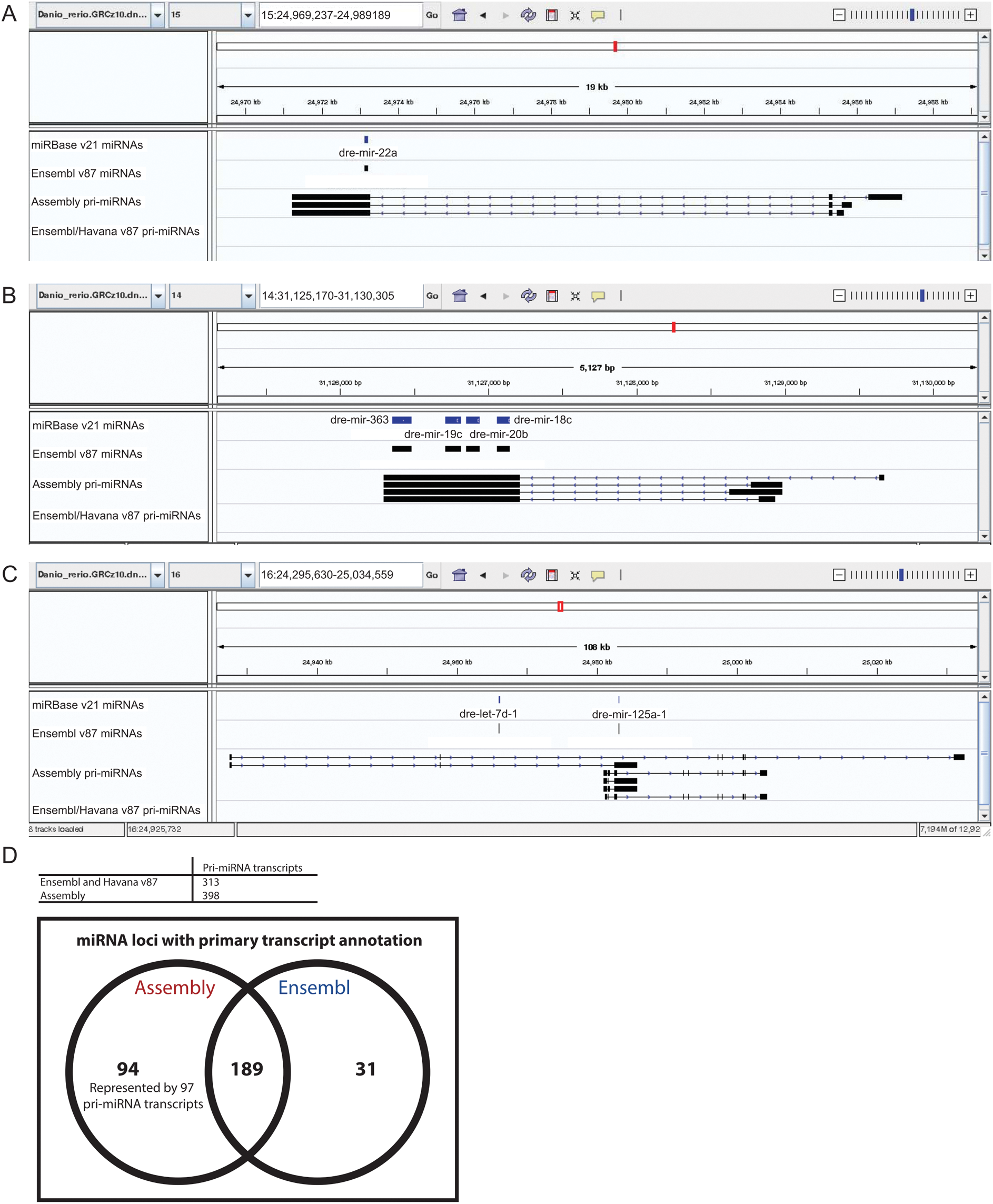
Annotation of primary miRNA transcripts. (A-C) A selection of miRNA loci for which pri-miRNA annotation is not available in Ensembl and Havana v87. IGV screen shots showing pri-miRNA transcripts overlapping mir-22a (A), mir-363, mir-19c, mir-20b and mir-18c (B), and mir-125 (C). Tracks shown are miRNA annotations from miRBase and Ensembl, newly assembled pri-miRNA transcripts and Ensembl pri-miRNAs. (D) Table and Venn diagram showing the numbers of assembled pri-miRNA transcripts and the overlap with of Ensembl miRNAs with an annotated pri-miRNA.

## Discussion

We have produced a high quality map of zebrafish mRNA expression during development. It represents the first detailed temporal mRNA reference in zebrafish and enhances our understanding of the transcriptional diversity underlying vertebrate development.

Ordering the gene set by the stage at which they have their maximum expression gives a detailed overview of development, showing the molecular correlates of morphological changes within the embryo. This arrangement of the dataset reveals that there is a much larger fraction of unnamed genes in the stages from dome to 75% epiboly. We find that this peak in unnamed genes during gastrulation is not driven by an enrichment for teleost or zebrafish-specific genes, as the proportion of unnamed genes with mammalian orthologues is at the same level during these stages as during the rest of development. This implies that there is a large number of genes involved in vertebrate gastrulation that have yet to be characterised.

Having a large number of stages and replicates allows us to cluster genes by their expression profile across all of development using a graph-based method. This splits the assayed genes into 252 different expression profiles containing at least 5 genes each (10,363 of the genes are assigned to clusters). GO/ZFA enrichment on these clusters allows us to associate unnamed genes with known processes.

One intriguing set of clusters contains an over-representation of genes on the long arm of chromosome 4, a region considered to be constitutive heterochromatin (Howe et al., 2013b). The vast majority of these genes encode proteins with zinc finger domains. The clusters show expression profiles that rise sharply around the time of zygotic genome activation (ZGA) and decrease at the latest after 75% epiboly. Given this, it is tempting to speculate that these genes have a role during the activation of the zygotic genome. In *Drosophila* the zinc finger gene Zelda (zld) is involved in zygotic genome activation by early and widespread binding to promoters and enhancers ahead of their genes’ activation during the maternal-zygotic transition (Harrison et al., 2010; Harrison et al., 2011; Liang et al., 2008). In zebrafish, *pou5f3*, *nanog* and *soxB* genes have been implicated in activating specific genes during ZGA (Lee et al., 2013; Leichsenring et al., 2013), but no general factor comparable to Zelda has been identified yet. It is possible that the role of the ZnF protein family genes on chromosome 4 is to set up or maintain open chromatin in a genome-wide way to allow for the rapid activation of the first zygotic genes. This is supported by the finding that these clusters also contain four unnamed genes, also found on chromosome 4, that are predicted to be H3K36 methyltransferases (ENSDARG00000104681, ENSDARG00000091062, ENSDARG00000076160, ENSDARG00000103283). H3K36 methylation is generally, albeit not exclusively, associated with active euchromatin (Wagner and Carpenter, 2012). Another option is that they could be responsible for negatively regulating areas of open chromatin to stop inappropriate activation of transposable elements during ZGA. In mammals, suppression of retrotransposons is mediated by the rapidly evolving KRAB zinc finger genes (Jacobs et al., 2014). KRAB zinc finger genes are exclusive to mammals, making the chromosome 4 ZnF genes a possible functional equivalent.

Also, as would be expected given their expression profiles, within these four clusters are a large number of well-known developmental regulators such as members of the Tgf-beta (Bmp and Nodal) and Wnt signalling pathways, as well as transcription factors such as *gata5* & *6*, *sox32*, *vox*, *ved*, *vent* and both *Brachyury* homologues.

Correlation analysis of paralogues stemming from a teleost-specific genome duplication shows that while there is the expected skew towards positive temporal expression correlation, a significant proportion of paralogues have diverged in their expression pattern as demonstrated by low or negative correlation coefficients. This is confirmed by an analysis of the expression annotations provided by ZFIN. There are caveats to this analysis, not least that the annotation is far from complete and, therefore, annotations may not overlap simply because a certain stage has been investigated for one paralogue, but not the other. It will also be the case that some genes have expression patterns that are diverged spatially, but not temporally. Thus the number of genes whose expression annotations do not overlap is an estimate, but it does suggest that the phenomenon of divergence is widespread among paralogues, something that might be expected to be the case since there needs to be some form of selective pressure on retaining paralogous pairs.

A further layer of complexity is added by differential exon and 3′ end use. The dataset uncovers new splice isoforms which, when viewed in Ensembl against the Ensembl genebuild, will refines the current gene annotation and ties splice isoforms to their temporal expression patterns. This is of particular relevance to the ongoing debate surrounding reverse genetics approaches in zebrafish (Kok et al., 2015). One possible explanation for phenotypic discrepancies between morpholino antisense oligo knockdown, code disrupting point mutations and CRISPR/Cas9 or TALEN-mediated larger deletions is that mutations affect potentially non-critical exons that are only present in a subset of transcripts of a given gene. An improved annotation that gives temporal resolution of exon usage will be of tremendous value for the identification of critical exons.

Using the 3′ end sequencing method DeTCT, we have catalogued the differential use of alternative 3′ ends in a filtered set of transcripts. In accordance with previous work on a subset of stages (Collins et al., 2012; Sheppard et al., 2013; Ulitsky et al., 2012) we find that there is differential use of not only 3′ UTRs but also coding sequence across development. We show that *pdlim5b* is expressed as several transcripts, with a single maternal isoform and multiple zygotic ones producing different proteins. Incorporation of the data into an updated gene set will systematically identify these and refines our understanding of gene regulation. Furthermore, it provides scope for temporally resolved analysis using transcript-specific knockout targeting strategies.

An important role of 3′ UTRs is in post-transcriptional regulation of their transcripts. The 3′ UTRs contain binding sites for miRNAs which silence transcripts through translational repression or mRNA destabilisation (Bartel, 2009). However, the primary transcripts of miRNAs are often poorly annotated. Using two miRNA processing mutants (*dgcr8* and *drosha*), we have produced an improved pri-miRNA assembly with an increase of 42% in the number of Ensembl miRNAs with an associated primary transcript. In agreement with previous studies (Chang et al., 2015; Gaeta et al., 2017), the assembly shows that miRNAs are processed from primary transcripts with multiple isoforms with potentially multiple transcription start sites (Figure 7). This diversity of transcripts means that the expression of miRNAs in relatively close proximity (e.g. dre-let-7d-1 and dre-mir-125a-1) could be regulated at multiple levels (Figure 7C). In some transcripts both let-7d-1 and mir-125a-1 are both intronic and in some mir-125a-1 is exonic. There are also transcripts with alternate promoters which exclude the let-7d-1 locus entirely. These features provide significant flexibility to the transcriptional regulation of miRNAs. It is possible that a proportion of the assembled constructs are fragments. Clear definition of the 5’ and 3’ termini will require further work and ultimately experimental validation by an alternative method such as 5′ rapid amplification of cDNA ends.

A major aim for this work was to create an accessible reference of normal mRNA expression during zebrafish development. The primary sequence is available from the European Nucleotide Archive (ENA, http://www.ebi.ac.uk/ena), but importantly has also been interpreted for immediate user-friendly access. Expression Atlas provides relative expression levels while Ensembl displays alternative genomic transcript structures alongside the current gene annotation. These pre-computed analyses allow an in-depth examination of the data on a gene by gene basis. This makes it easy for researchers to benefit from the data and provides a direct link to the wealth of information present in genomic databases.

## Materials and Methods

### Animal Care

Zebrafish were maintained in accordance with UK Home Office regulations, UK Animals (Scientific Procedures) Act 1986, under project licence 70/7606, which was reviewed by the Wellcome Trust Sanger Institute Ethical Review Committee.

### Sample collection

Wild-type HLF strain zebrafish (*Danio rerio*) were maintained at 28.5 °C on a 14h light/10h dark cycle. At the time of mating, breeding males and females were separated overnight before letting them spawn naturally in the morning to allow for synchronisation of developmental stages. Fertilised eggs were grown at 28°C and staged using previously defined criteria (Kimmel et al., 1995). Samples from 18 different developmental stages from 1-cell to 5 days post fertilisation were collected by snap freezing the embryos in dry ice. Details of sample names and accession numbers are provided in Supplemental Data File 1.

### RNA Extraction

RNA was extracted from embryos by mechanical lysis in RLT buffer (Qiagen) containing 1 μl of 14.3M beta mercaptoethanol (Sigma). The lysate was combined with 1.8 volumes of Agencourt RNAClean XP (Beckman Coulter) beads and allowed to bind for 10 minutes. The plate was applied to a plate magnet (Invitrogen) until the solution cleared and the supernatant was removed without disturbing the beads. This was followed by washing the beads three times with 70% ethanol. After the last wash, the pellet was allowed to air dry for 10 mins and then resuspended in 50 μl of RNAse-free water. RNA was eluted from the beads by applying the plate to the magnetic rack. RNA was quantified using Quant-IT RNA assay (Invitrogen) for samples 24 hours post-fertilisation and older.

### RNA Sequencing

Total RNA from 12 embryos was pooled and DNase treated for 20 mins at 37°C followed by addition of 1 μl 0.5M EDTA and inactivation at 75°C for 10 mins to remove residual DNA. RNA was then cleaned using 2 volumes of Agencourt RNAClean XP (Beckman Coulter) beads under the standard protocol. Five replicate libraries were made for each stage using 700 ng total RNA and ERCC spike mix 2 (Ambion). Strand-specific RNA-seq libraries containing unique index sequences in the adapter were generated simultaneously following the dUTP method. Libraries were pooled and sequenced on Illumina HiSeq 2500 in 100bp paired-end mode. Sequence data were deposited in ENA under accession ERP014517.

FASTQ files were aligned to the GRCz10 reference genome using tophat (v2.0.13, options: --library-type fr-firststrand). Counts for genes were produced using htseq-count (v0.6.0 options: --stranded=reverse) with the Ensembl v85 annotation as a reference. The processed count data are in Supplemental Data File 2.

### DeTCT

DeTCT libraries were generated, sequenced and analysed as described previously (Collins et al., 2015). The resulting genomic regions and putative 3′ ends were filtered stringently using DeTCT’s filter_output script (https://github.com/iansealy/DETCT/blob/master/script/filter_output.pl) in its --stricter mode. This mode removes 3′ ends more than 5000 bases downstream of existing protein-coding annotation (Ensembl v86) or more than 50 bases from existing non-coding annotation (antisense, lincRNA, misc_RNA and processed_transcript biotypes). 3′ ends are also removed if nearby sequence is enriched for As, if within 14 bp of an N, if within a simple repeat (repeats annotated by dust or trf) or a transposon, or if not nearby a primary hexamer. Regions not associated with 3′ ends are also removed. Sequence data were deposited in ENA under accessions ERP006948 and ERP013756. The processed count data is available to download at https://doi.org/10.6084/m9.figshare.4622311.

## Analysis

### ZFA enrichment

ZFA enrichment was done using Ontologizer (http://ontologizer.de). The ZFA ontology was download from http://www.berkeleybop.org/ontologies/zfa.obo and mappings between ZFIN gene IDs and ZFA IDs was downloaded from ZFIN (http://zfin.org/downloads/wildtype-expression_fish.txt and http://zfin.org/downloads/phenoGeneCleanData_fish.txt). Pvalues were adjusted for testing of both multiple terms and multiple clusters using the Bonferroni correction.

### GO enrichment

GO enrichment was done using the R topGO package (Alexa and Rahnenfuhrer, 2016). The mapping between Ensembl IDs and GO terms was retrieved from the Ensembl database using a custom perl script (get_ensembl_go_terms.pl) from the topgo-wrapper repository (https://github.com/iansealy/topgo-wrapper).

### Plots

The code to reproduce all the plots in this paper can be downloaded at github (https://github.com/richysix/zfs-devstages).

### BioLayout

BioLayout *Express*^3D^ is available to download for free (http://www.biolayout.org/downloads/). The input expression file is provided as Supplemental Data File 6. The processed layout file of the network displayed in Figure 3B is available at https://doi.org/10.6084/m9.figshare.4622293.

### pri-miRNA assembly

For each mutant allele – sa191, *drosha* (*rnasen*); sa223, *dgcr8* (*pasha*) – we produced and sequenced stranded RNA-seq libraries from homozygous, heterozygous and wild-type embryos (5 replicates of each). Sequence data were deposited in ENA under accession ERP013690. To identify new pri-miRNAs, FASTQ files were aligned to the zebrafish genome (GRCz10) using STAR (v2.5.1a, options: --alignSJoverhangMin 8 --alignSJDBoverhangMin 1 --outFilterMismatchNmax 999 -- alignIntronMin 20 --alignIntronMax 1000000 --alignMatesGapMax 1000000 -- readFilesCommand zcat --outFilterIntronMotifs RemoveNoncanonical --outFilterType BySJout --outFilterMultimapNmax 10 --sjdbOverhang 74). Ensembl v86 zebrafish annotation was provided as a junction file. Aligned reads were sorted, indexed and mate pairings were fixed using Samtools (v1.3.1). Duplicate alignments were removed with Picard (v2.7.1) (REMOVE_DUPLICATES=true READ_NAME_REGEX=null). Alignments for the homozygous, heterozygous and wild-type samples for both the *drosha* and *dgcr8* lines were merged into two line-specific datasets with Samtools. Each set of merged alignments was then used to assemble a transcriptome with Cufflinks (v2.2.1) (--library-type=fr-firststrand). Ensembl (v86) annotation stripped of rows corresponding to ‘gene’ features was provided as a backbone for the assembly. The *drosha* and *dgcr8* assemblies were then merged with Cuffmerge.

Ensembl miRNA annotation (v87) (“miRNA” biotype) was used to define miRNA loci. For the purpose of this analysis we defined pri-miRNAs as any spliced transcript overlapping at least one of these loci. Ensembl and Havana (v87) annotation was selected as the baseline for novel pri-miRNA prediction. UCSC utilities gtfToGenePred and genePredToBed were used to convert from GTF to BED format. Pri-miRNAs were identified by removing single exon transcripts from the merged assembly and Ensembl annotations and comparing the remainder to annotated miRNA loci using bedtools (v2.24.0) intersect (-wa –s). Finally, assembled pri-miRNA transcripts were filtered to remove those for which either the first or last exon was less than 20nt in length, those which contain an intron greater than 100,000nt, or those which contain an intron that isn’t flanked by the GT-AG or GC-AG motifs (Breathnach et al., 1978).

## Acknowledgements

This work was funded by the Wellcome Trust Sanger Institute (grant number WT098051). M. Davis was funded by a MRC Methodology Research Fellowship (MR/L012367/1). Ensembl, Expression Atlas and A. Enright are funded by EMBL core funding. We would like to thank the Wellcome Trust Sanger Institute sequencing pipelines for performing sequencing and the staff of the Research Support Facility for zebrafish care. We also thank Samantha Carruthers, Catherine Scahill and Nicole Staudt for critical reading of the manuscript.

## Supplemental Data Files

Supplemental Data File 1 – Sample Information (.tsv)

Supplemental Data File 2 – RNA-seq count data (.tsv)

Supplemental Data File 3 – Max stage cluster assignments (.tsv)

Supplemental Data File 4 – Max stage cluster ZFA enrichment (.tsv)

Supplemental Data File 5 – Max stage cluster GO enrichment (.tsv)

Supplemental Data File 6 – BioLayout input file (.expression)

Supplemental Data File 7 – BioLayout cluster information (.tsv)

Supplemental Data File 8 – BioLayout cluster ZFA enrichment (.zip)

Supplemental Data File 9 – BioLayout cluster GO enrichment (.tsv)

Supplemental Data File 10 – Regional expression information (.tsv)

Supplemental Data File 11 – Assembled pri-miRNA transcripts (.bed)

Supplemental Data File 12 – Table of overlap of miRNAs (.xlsx)

